# A malaria parasite phospholipid flippase safeguards midgut traversal of ookinetes for mosquito transmission

**DOI:** 10.1101/2021.03.14.435214

**Authors:** Zhenke Yang, Yang Shi, Huiting Cui, Shuzhen Yang, Han Gao, Jing Yuan

## Abstract

Mosquito midgut epithelium traversal is an essential component of transmission of malaria parasites. Phospholipid flippases are eukaryotic type IV ATPases (P4-ATPases), which in association with CDC50 cofactors, translocate phospholipids across lipid bilayers to maintain the membrane asymmetry. In this study, we investigated the function of a putative P4-ATPase, ATP7, from the rodent malaria parasite *P. yoelii*. Disruption of ATP7 results in block of parasite infection of mosquitoes. ATP7 is localized on the ookinete plasma membrane. While ATP7-depleted ookinetes are motile and capable of invading the midgut, they are quickly eliminated within the epithelial cells by a process that is independent from the mosquito complement-like immunity. ATP7 colocalizes and interacts with the flippase co-factor CDC50C. Importantly, depletion of CDC50C phenocopies ATP7 deficiency. ATP7-depleted ookinetes fail to translocate phosphatidylcholine (PC) across the plasma membrane, resulting in PC exposure at the ookinete surface. Lastly, ookinete microinjection into the mosquito hemocoel reverses the ATP7 deficiency phenotype. Our study identifies *Plasmodium* flippase as a novel mechanism of parasite survival in the midgut epithelium that is required for mosquito transmission.

## Introduction

Malaria is among the deadliest parasitic diseases and numbered an estimated 219 million cases and 435,000 deaths in humans in 2018 [1]. Malaria is caused by protozoan parasites of the genus *Plasmodium* that are transmitted between humans by female anopheline mosquitoes. Mosquito acquisition of *Plasmodium* starts when a mosquito ingests an infected human blood containing gametocytes. Once in the mosquito midgut lumen, female and male gametocytes immediately transform into extracellular gametes. Fertilization of gametes leads to a spherical zygote that within 12-20 hours further differentiates into a crescent-shaped motile ookinete. The ookinetes invade and traverse the single-layer midgut epithelium before reaching the sub-epithelial basal space where they transform into stationary oocysts. Thousands of sporozoites form within the oocyst over a period of 10-18 days. Upon release into the hemocoel, sporozoites migrate to mosquito salivary glands from where they are released when the mosquito bites a new host [2].

Upon midgut traversal, parasite numbers decrease dramaticaly due to the complement-like immune responses mediated by mosquito TEP1 (the homolog of human complement factor C3), and two TEP1-interacting proteins LRIM1 and APL1C [3–9]. Mosquito hemocytes containing phagocytic cells have also been implicated in the complement-like immunity against *Plasmodium* [10]. In addition, ookinete invasion triggers apoptosis of the traversed cell, causing extrusion and clearance of the epithelium into the lumen [11]. *Plasmodium* has also evolved mechanisms to avoid host attack. Parasite surface proteins have been implicated in conferring resistance to anti-*Plasmodium* immunity. A variety of ookinete surface proteins interact with the mosquito host promoting parasite infection, survival, and transmission [12]. The ookinete surface protein P47, a member of six-cysteine protein family [13, 14], suppresses mosquito JNK signaling in the invaded midgut cell, inhibiting ookinete nitration and destruction by complement-like reactions [15–17]. Another parasite surface protein PIMMS43 was recently shown to protect ookinetes from the mosquito complement-like immunity [18]. It is generally accepted that both P47 and PIMMS43 play critical roles to enable ookinete evade mosquito complement-like attack upon arriving at the sub-epithelial basal space. However, mosquito anti-*Plasmodium* immunity and ookinete defenses remain incompletely understood.

Asymmetric distribution of lipid molecules across the two leaflets of biological membranes is an important feature of eukaryotic cells. To establish asymmetry, eukaryotic organisms ranging from yeast to human encode a specific type of membrane transporters known as flippases that translocate phospholipids to the cytosolic side of membranes in a reaction driven by ATP [19]. These flippases belong to the P4-ATPase subfamily of P-type ATPases [19] and CDC50 proteins act as essential P4-ATPase co-factors [19]. Primate malaria parasites encode four putative canonical P4-ATPases while rodent malaria parasites encode three. However, the precise roles of these P4-ATPases in malaria parasite infection, development, and transmission have not been established.

Recently, we performed a CRISPR/Cas9-based screen of a large number of *Plasmodium yoelii* membrane protein-encoding genes to search for genes critical for development in the mosquito [20]. Notably, one gene (PY17X_0809500) was determined to be essential for *P. yoelii* parasite development in the mosquito. This gene encodes a putative P4-type ATPase protein (designated as ATP7 in this study) that is conserved among *Plasmodium* species. In this study, we performed in-depth *in vitro* and *in vivo* analyses of the *P. yoelii* P4-ATPase (PY17X_0809500) using gene disruption. We found an essential role for ATP7 and its co-factor CDC50C, in ookinete survival and evasion from mosquito midgut immunity.

## Results

### P4-ATPase ATP7 is specifically expressed in the mosquito stages of *Plasmodium*

The *P. yoelii* PY17X_0809500 gene encodes a 1764 amino acid protein with 10 predicted transmembrane helixes with extensive identity to key P4-ATPase subdomains (Fig. 1A). To investigate the temporal expression and subcellular localization of ATP7 in the parasite, we tagged endogenous ATP7 with a sextuple HA epitope (6HA) at the C-terminus in the *P. yoelii* 17XNL strain by homologous double cross-over using a CRISPR/Cas9-based approach [21, 22](Fig. S1). The resulting *atp7::6HA* parasite showed normal development throughout its life cycle. Immunoblot and immunofluorescence assay (IFA) showed that ATP7 was expressed in gametocytes, mosquito midgut oocysts, and salivary gland sporozoites, but was not detected in asexual blood stage parasites (Fig. 1B and C). Interestingly, ATP7 was detected in the cytoplasm of gametocytes and oocysts, but displayed a peripheral localization in sporozoites (Fig. 1C). Co-staining of the *atp7::6HA* gametocytes with α-tubulin (male gametocyte specific) and HA antibodies showed that ATP7 was expressed only in female gametocytes (Fig. 1D). During zygote to ookinete differentiation *in vitro*, ATP7 was distributed in both the cytoplasm and the cell periphery from zygote to retort, but was mostly localized to the cell periphery of mature ookinetes (Fig. 1E). Furthermore, we tagged the endogenous ATP7 with a quadruple Myc epitope (4Myc) (Fig. S1) and obtained similar results in the *atp7::4Myc* parasites (Fig. 1F and G). In summary, ATP7 is specifically expressed in female gametocytes and then throughout parasite development in the mosquito.

**Figure 1.**
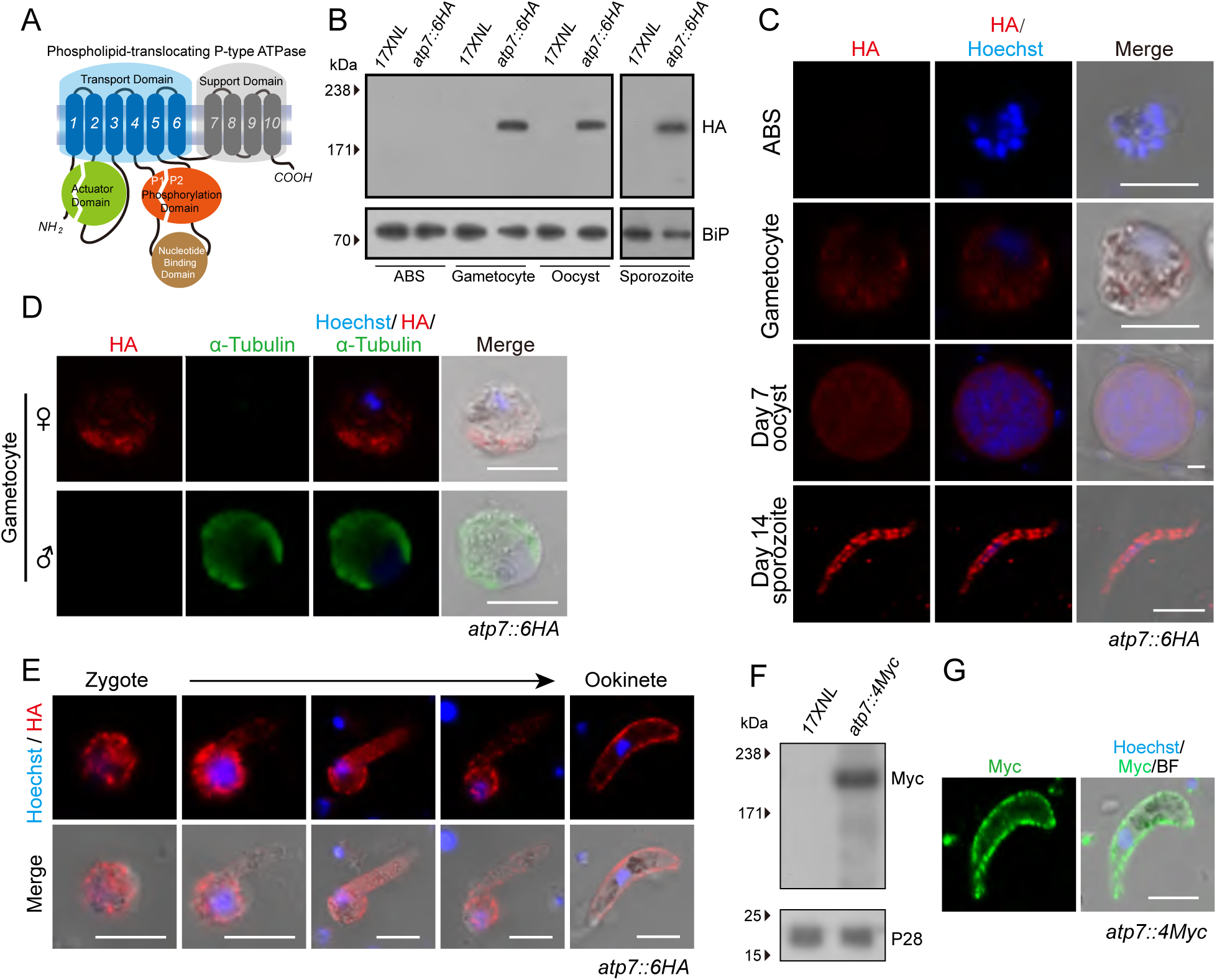
The P4-ATPase ATP7 is expressed in *Plasmodium* mosquito stages. **A.** Predicted protein topology of *P. yoelii* P4-type ATPase ATP7 (PY17X_0809500). Transmembrane helixes 1-10, actuator domain (green), phosphorylation domain (orange), and nucleotide binding domain (brown) are indicated. **B.** Immunoblot of ATP7 at asexual blood stages (ABS), gametocytes, midgut oocysts (7 days post blood feeding) and salivary gland sporozoites (14 days) of the 17XNL (wildtype parasite) and the tagged parasite *atp7::6HA*. ER protein BiP was used as a loading control. **C.** IFA of ATP7 expression in ABS, gametocytes, midgut oocysts, and salivary gland sporozoites of the *atp7::6HA* parasite. Nuclei are labeled with Hoechst 33342. **D.** Co-staining *atp7::6HA* gametocytes with HA and male-gametocyte-specific *α*-tubulin II antibodies. **E.** IFA for ATP7 expression dynamics during *in vitro* zygote to ookinete differentiation of the *atp7::6HA* parasite. **F.** Immunoblot of ATP7 in the ookinetes of the tagged parasite *atp7::4Myc*. P28 protein was used as a loading control. **G.** IFA of ATP7 expression in ookinetes of another tagged parasite *atp7::4Myc*. Scale bar = 5 μm for all images. Parasites were permeabilized by Triton X-100 in IFA analysis. All experiments in this figure were independently repeated three times with similar results, and the data are shown from one representative experiment.

### ATP7 is essential for transmission of malaria parasites to the mosquito

To investigate the function of ATP7 in the parasite life cycle, we generated Cas9 constructs to disrupt the *atp7* gene in *P. yoelii* (strain 17XNL) by homologous double cross-over (Fig. S1). This gene has a single exon with a 5295 bp coding region. We successfully obtained a mutant clone *Δatp7c* bearing a 1 kb deletion of the C-terminal coding region (Fig. 2A and Fig. S1). The *Δatp7c* clone exhibited normal development of asexual blood stages as well as gametocytes in the mice (Fig. 2D and E).

**Figure 2.**
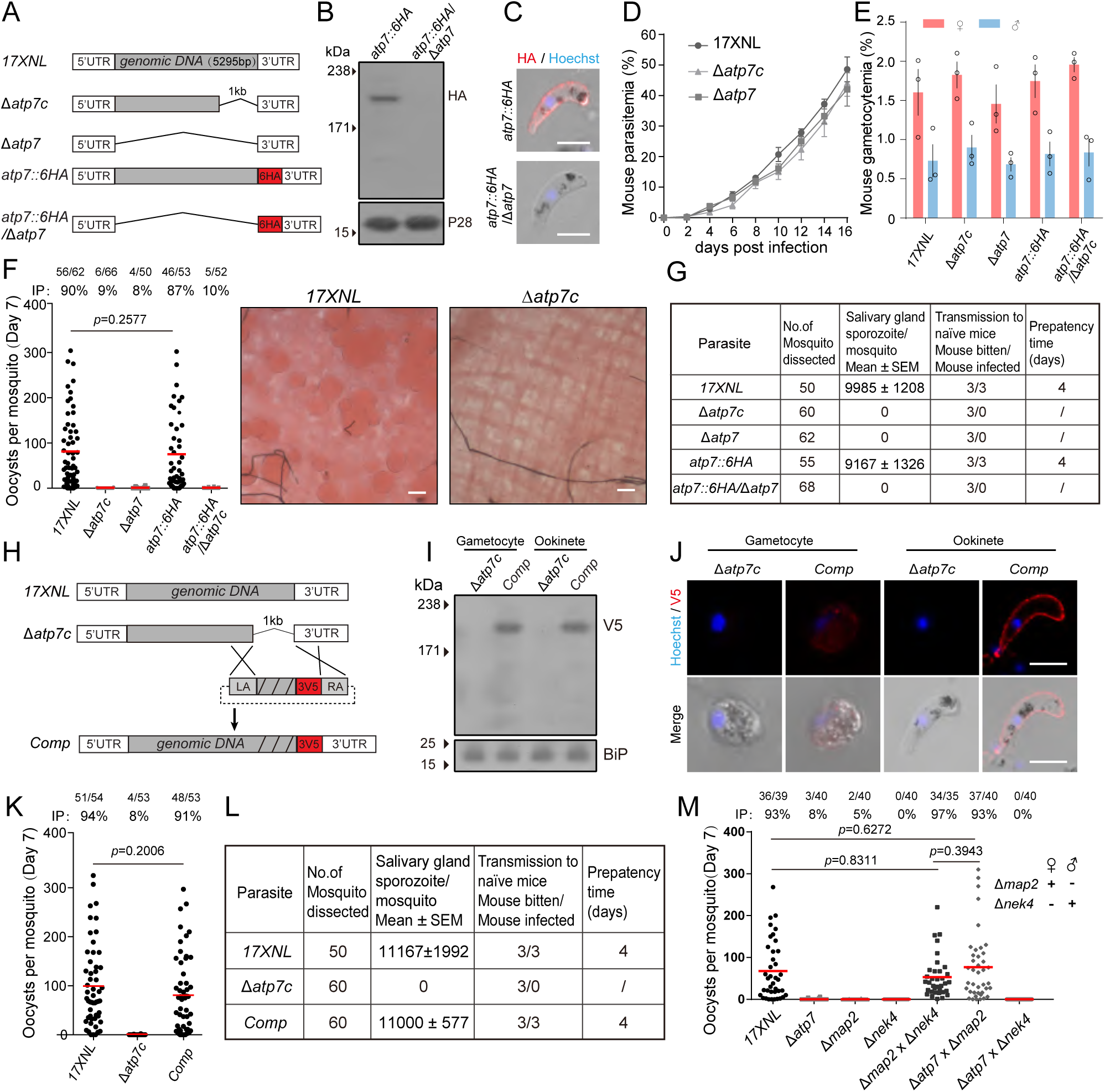
ATP7 is dispensable for asexual stages but essential for mosquito stages of parasites. **A.** Diagram showing genetic disruption of the *atp7* gene in the 17XNL and *atp7::6HA* parasites background. C-terminus (1 kb) or full length (5.3 kb) of the coding sequence was deleted using CRISPR/Cas9 method (see Figure S1). **B.** Immunoblot of ATP7 expression in *atp7::6HA* and *atp7::6HA*/Δ*atp7* ookinetes. P28 was used as a loading control. **C.** IFA of ATP7 in *atp7::6HA* and *atp7::6HA*/Δ*atp7* ookinetes. Ookinetes were permeabilized by Triton X-100. Scale bar = 5 μm. **D.** Asexual blood stage proliferation of 17XNL, Δ*atp7*, and Δ*atp7c* parasites. **E.** Gametocyte formation of the 17XNL, Δ*atp7*, Δ*atp7c*, *atp7::6HA*, and *atp7::6HA*/Δ*atp7* parasites in mice. **F.** Midgut oocyst formation in mosquito 7 days post blood feeding. Right panel shows midguts stained with 0.5% mercurochrome. Scale bar = 50 μm. **G.** Salivary gland sporozoite counts and mouse infectivity. For each group, 20 mosquitoes were fed on one mouse and the prepatent time was determined. **H.** Diagram of Δ*atp7c* mutant complementation using CRISPR/Cas9. A triple V5 (3V5, red) epitope was fused to the gene. **I.** Immunoblot confirming the expression of 3V5-tagged ATP7 in gametocytes and ookinetes of the complemented parasite (*Comp*). BiP was used as a loading control. **J.** IFA confirming 3V5-tagged ATP7 expression in gametocytes and ookinetes of the complemented parasite. Scale bar = 5 μm. **K.** Midgut oocyst numbers in mosquitoes infected with the complemented parasite. **L.** Salivary gland sporozoite counts and mouse infectivity of the complemented parasite. **M.** Midgut oocyst formation in mosquitoes infected with parasites, including 17XNL, △*atp7*, △*map2*, or △*nek4* parasite alone as well as parasite combinations of △*map*2*/*△*nek*4, △*atp7/*△*map*2, and △*atp7/*△*nek*4. △*nek*4 and △*map*2 are strains that do not produce female and male gametes, respectively. Data are shown as mean ± SD in **D** and **E**; in **F**, **K**, and **M**, the nubers on the top are the number of mosquito carrying oocysts / the number of mosquito dissected; IP: mosquito infection prevalence expressed as percentages; red horizontal lines show mean oocyst number values; and Mann–Whitney test applied. Experiments in this figure were independently repeated three times except two times in **M**, and the data are shown from one representative experiment.

To evaluate the role of ATP7 in parasite development in the mosquito, *An. stephensi* mosquitoes were fed on infected mice. The mosquitoes were dissected and the formed oocysts in the midguts were counted after mercurochrome staining. We found that *Δatp7c* parasites did not produce any oocyst on day 7 post-infection (pi) (Fig. 2F), indicating failure of mosquito transmission in the absence of this gene. Consistent with this finding, no sporozoites were detected in the salivary glands on day 14 pi and no transmission from mosquitoes to mice was observed (Fig. 2G). To confirm that the defects were indeed caused by ATP7 deletion, we re-introduced the deleted 1.0 kb sequence fused with a triple V5 epitope (3V5) coding sequence, back into the *atp7* locus of the *Δatp7c* parasite strain (Fig. 2H and Fig. S1). Expression of V5-tagged ATP7 was detected in gametocytes and ookinetes from the complemented parasites using immunoblot (Fig. 2I) and IFA (Fig. 2J). In line with the localization of the endogenous ATP7 (Fig. 1E), the ATP7::3V5 fusion protein also exhibited a peripheral localization in the complemented ookinetes (Fig. 2J). Notably, oocyst and sporozoite formation, as well as mouse infectivity, were all restored (Fig. 2K and L). To further confirm that the defects of the *Δatp7c* strain resulted directly from *atp7* deficiency, we generated two additional parasite mutants by deleting the entire coding region of the *atp7* gene in the WT and *atp7::6HA* strains respectively (Fig. 2A-C). Consistent with the findings with the *Δatp7c* mutant, these *Δatp7* and *atp7::6HA*/*Δatp7* mutants were also able to generate gametocytes in mice (Fig. 2E) but did not produce oocysts or sporozoites in mosquitoes (Fig. 2F and G). To confirm the female-inheritance of ATP7 function, we performed a genetic cross of the Δ*atp7* with either Δ*map2* (male gamete-deficient) or Δ*nek4* (female gamete-deficient) parasites [20]. Normal numbers of midgut oocysts were observed in mosquitoes on day 7 pi in the Δ*atp7*×Δ*map2* but not the Δ*atp7*×Δ*nek4* cross (Fig. 2M), in agreement with the specific expression of ATP7 in female gametocytes (Fig. 1D).

### ATP7-deficient *P. yoelii* produce ookinetes but not early oocysts

In the lumen of the mosquito midgut, malaria parasites undergo a series of distinct stage transformations (gametocyte to gamete to zygote and to ookinete) before traversing the midgut epithelium to initiate differentiation into oocysts in the sub-epithelial basal space. We performed experiments to investigate which stages are affected by ATP7 deficiency. The *Δatp7* parasites produced normal male and female gametes *in vitro* (Fig. 3A). Similarly, zygote to ookinete differentiation *in vitro* (Fig. 3B and C) and *in vivo* were normal in the mutant parasites compared to controls (Fig. 3D and E). Scanning electron microscopy (SEM) showed crescent-shaped *Δatp7* ookinetes with an intact apex indistinguishable from that of WT ookinetes (Fig. 3F). Since gliding motility is a prerequisite for midgut invasion by ookinetes, we assessed their gliding capability using an *in vitro* Matrigel-based assay and observed no appreciable difference in gliding speed between the WT and *Δatp7* (Fig. 3G). In contrast, ookinetes lacking the *ctrp* gene, which is essential for ookinete gliding [23, 24], completely lost motility (Fig. 3G). Moreover, we tested whether ATP7 regulates microneme secretion, which is essential for ookinete gliding and invasion [24–26]. Comparable levels of the secreted proteins from the microneme including CTRP, chitinase, and WARP were detected in the supernatants of WT and mutant ookinete cultures (Fig. S2A), suggesting normal microneme secretion. Additionally, transmission electron microscopy (TEM) revealed intact micronemes, apical ring, and apical collar in the apex of *Δatp7* ookinetes (Fig. S2B and C). Further experiments showed that WT and the ATP7-depleted ookinetes exhibit similar cell morphology and survival under different stress conditions *in vitro* (Fig. S2D-I). Together, the ATP7-depleted parasite is able to develop mature and fully motile ookinetes.

**Figure 3.**
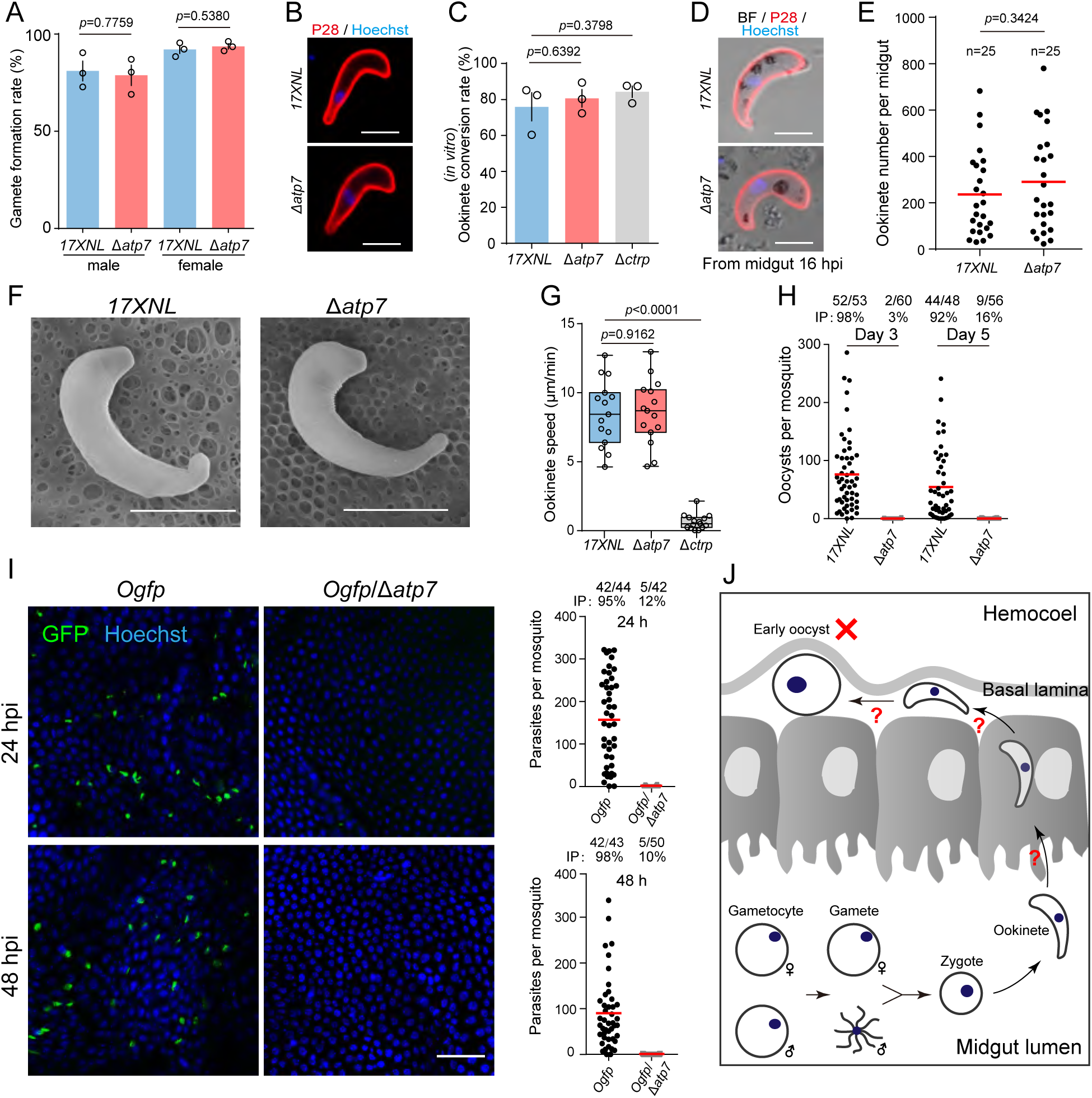
ATP7 deficient parasites develop ookinetes but no early oocysts. **A.** Gametocyte activation to gamete *in vitro*. Male and female gamete formation rates are the percentage of male gametocytes showing exflagellation and the percentage of female gametocytes showing P28 expression after XA stimulation. **B.** IFA of P28 in 17XNL and Δ*atp7* ookinetes *in vitro*. P28 is an ookinete plasma membrane protein. **C.** Quantification of ookinete formation in **B**. Ookinete conversion rate is the number of ookinete per 100 female gametocytes. Means ± SEM from 3 independent tests. **D.** IFA of P28 in ookinetes from infected mosquito midguts 16 h pi. **E.** Quantification of the ookinetes in **D**. n is the number of mosquitoes dissected. **F.** Ookinete images by scanning electron microscope (SEM). **G.** Ookinete gliding motility measured using the *in vitro* Matrigel-based assay. **H.** Midgut oocysts numbers at 3 and 5 day pi. **I.** Fluorescence microscopy observation of ookinetes and early oocysts in mosquito midguts infected with the reporter strains *Ogfp* and *Ogfp*/*Δatp7* at 24 and 48 h pi. Right panels show the quantification of parasites. **J.** Model for ATP7 deficiency in mosquito transmission. Scale bar is 5 μm in **B**, **D**, and **F**, and 50 μm in **I**. Data is showed as mean ± SD in **A** and **C**. In **H** and **I**, the numbers on the top are the number of mosquito carrying oocysts / the number of mosquito dissected; IP: mosquito infection prevalence expressed as percentages. Red horizontal lines show mean oocyst number values. Two-tailed unpaired Student’s t test in **A**, **C**, **E** and **G**. All experiments were repeated three times, and the data are shown from one representative experiment.

We next investigated the development of early oocysts. Compared to WT, no *Δatp7* oocysts were detected in mosquito midguts on both day 3 and 5 pi (Fig. 3H). Unlike late stage oocysts, early stage oocysts (day 1-2 pi) cannot be easily visualized with mercurochrome staining. Since the *soap* gene is highly expressed in mature ookinetes and early oocysts [27], we engineered two parasite strains *Ogfp* and *OmCherry*, wherein the endogenous *soap* sequence is fused with the coding sequence of GFP and mCherry, respectively, at the C-terminus to track early oocysts (Fig. S3 and Fig. S1). A “ribosome skip” T2A peptide was inserted between the SOAP and GFP or mCherry to ensure the separate expression of GFP or mCherry and the endogenous SOAP. As expected, GFP and mCherry fluorescence was detected specifically in the mature ookinetes *in vitro* and in the ookinetes and early oocysts *in vivo* (Fig. S3A and B). Disruption of *atp7* in the *Ogfp* parasites led to almost complete absence of GFP^+^ cells in mosquito midguts on both 24 and 48 hour pi (Fig. 3I), clearly indicating a lack of early oocysts (Fig. 3J).

### ATP7-depleted ookinetes are capable of invading mosquito midgut epithelium

Four possibilities may account for the fact that ATP7-depleted parasites develop to motile ookinetes but fail to form early oocysts in the midgut (Fig. 3J): 1. the ookinetes fail to invade midgut epithelium; 2. the ookinetes are developmentally arrested within the midgut epithelial cells; 3. the ookinetes are eliminated by the mosquito immune system; 4. the ookinetes fail to transform into early oocysts after traversal. To address the first possibility, we analyzed the transcriptional activation of the *An. stephensi* gene *AsSRPN6*, a hallmark of ookinete invasion [28], in the midguts of infected mosquitoes at 24 h pi. As expected, *AsSRPN6* mRNA remained minimally expressed in mosquitoes that fed on naïve mouse blood (control) as determined by qRT-PCR, and was significantly elevated in mosquitoes fed on blood carrying WT and *Δatp7* but not *Δctrp* parasites (Fig. S4A). Similar results were obtained when AsSRPN6 protein was assayed (Fig. S4B). Additionally, we examined tyrosine nitration and peroxidase activity, which are induced in epithelial cells invaded by ookinetes [29]. Tyrosine nitration was detected at 24 h pi in midguts infected with WT and *Δatp7* parasites, but not in control midguts or midguts infected with *Δctrp* parasites (Fig. S4C). Similarly, no differences were detected in peroxidase activity between midguts infected with WT and *Δatp7* parasites (Fig. S4D).

To further confirm successful epithelium invasion by the *atp7*-deficient ookinetes, we examined the location of ookinetes in midguts using confocal microscopy. Parasite-infected midguts were dissected at 18 h pi and visualized after staining with an antibody against P28. P28 is a plasma membrane protein of ookinetes and early oocysts [30]. On average, 87% fewer P28^+^ *atp7*-deficient parasites were detected compared with WT (Fig. S4E and F), recapitulating the results of *Ogfp*/*Δatp7* ookinetes (Fig. 3I). However, the percentage of *Δatp7* parasites (96%) localizing at the basal side is not notably different from that of the WT (97%) (Fig. S4F), indicating successful epithelium invasion of the *Δatp7* ookinetes. As was previously reported, no ookinetes were detected at the basal side in the *Δctrp* and *Δpplp5* mutants that are motility-defective and invasion-defective, respectively [24, 31] (Fig. S4E and F). Together these experiments show that ATPase7-depleted ookinetes are capable of invading the mosquito midgut epithelium (Fig. S4G).

### *Δatp7* ookinetes are eliminated during epithelium traversal

We next asked if the ATP7-deficient ookinetes are eliminated during midgut traversal. In parasite-infected midguts dissected at 24 h pi, the number of P28^+^ *Δatp7* ookinetes (mean number: 9) significantly decreased compared with the WT ookinetes (mean number: 112), although the mosquito infection level is comparable (infection prevalence is 90% for the WT and 86% for the *Δatp7*) (Fig. 4A). Notably, the *Δatp7* parasites in the midguts were morphologically aberrant. A total of 99% of the WT parasites had an intact shape (crescent or round) while 94% of the *Δatp7* mutants lost cell integrity and appeared to be lysed (Fig. 4B). In addition, ookinete nuclei were barely detected in *Δatp7* ookinetes in contrast with the intact nuclei observed in controls (Fig. 4B). Staining with antibody against GAP45, another pellicle protein in the parasite inner membrane complex (IMC), revealed similar defects (Fig. 4C). These results suggest drastic elimination of the ookinetes rather than developmental arrest within the epithelium. Moreover, deleting *atp7* in the *Ogfp* strain resulted in only 0.5% (2/375) of the GFP^+^ cells among those were labeled by P28 in comparison to 90% (530/587) in the controls (Fig. 4D-E). These results are consistent with the previous observations that GFP fluorescence rapidly fades away as the ookinetes are eliminated during midgut traversal, with P28 continuing to be on the cell surface [3, 15]. Together these results suggest that ATP7-depleted ookinetes are eliminated within the midgut epithelium (Fig. 4F).

**Figure 4.**
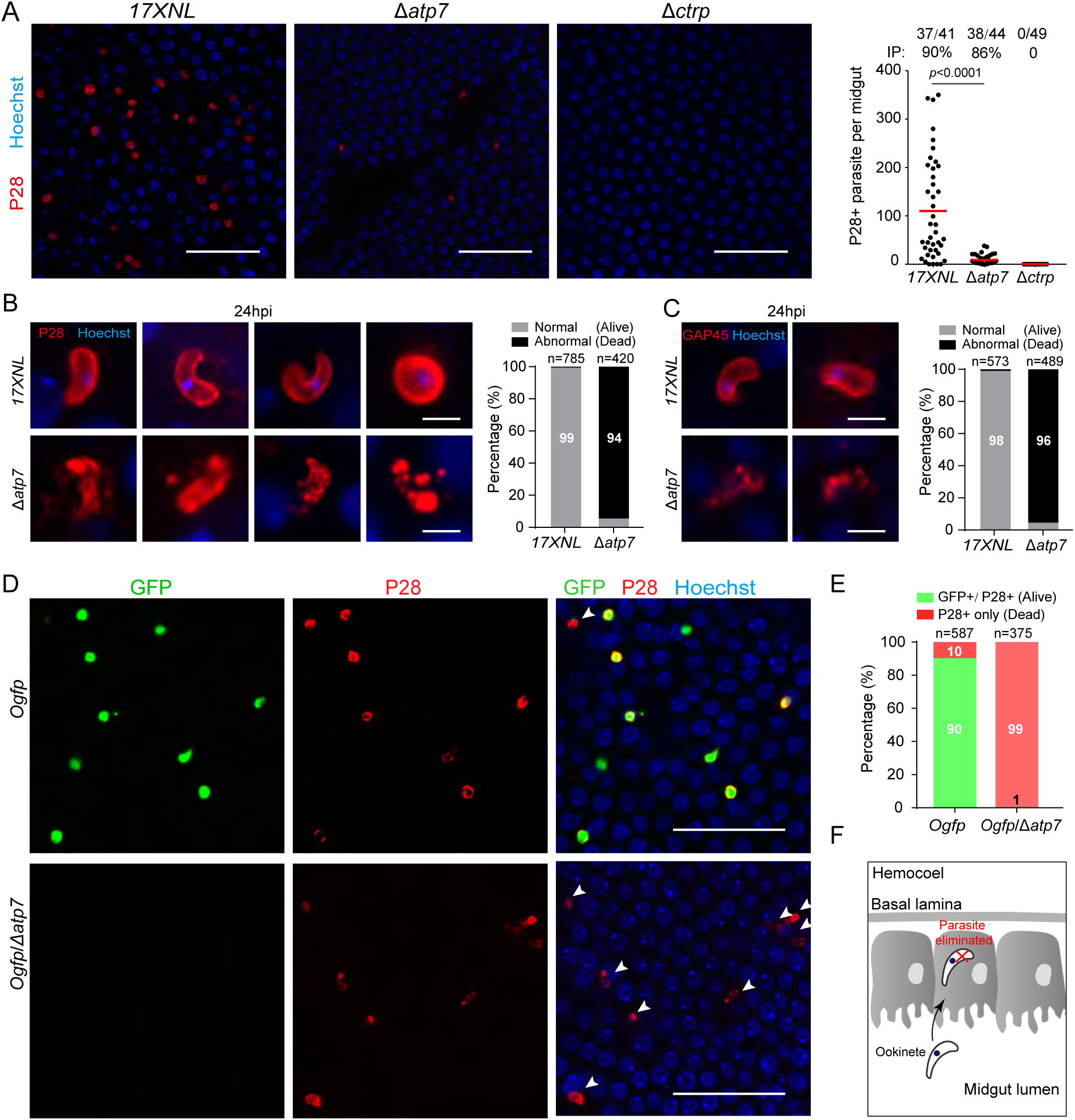
Δ*atp7* ookinetes are eliminated during midgut traversal. **A.** Parasite plasma membrane and nucleus staining in the infected midguts at 24 h pi. P28 is a plasma membrane protein of ookinete. Right panel is the quantification of the number of P28^+^ parasites per midgut. **B.** Enlarged images of P28^+^ parasites in **A**. Right panel shows the percentage of normal (alive) and abnormal (dead) parasites. n is the number of parasites counted. **C.** Parasite inner membrane complex and nucleus staining in the infected midguts at 24 h pi. GAP45 is an IMC protein of ookinete. Right panel shows the percentage of normal and abnormal parasites. n is the number of parasites counted. **D.** P28 and Hoechst 33342 staining of midguts infected with fluorescent reporter strains *Ogfp* and *Ogfp*/*Δatp7* at 24 h pi. White arrows indicate dead parasites showing P28 staining but no GFP fluorescence. **E.** Percentage of normal (alive) and abnormal (dead) parasites in **D**. n is the number of parasites counted. **F.** Diagram illustrating elimination of ATP7-depleted ookinetes within epithelium. Scale bar is 50 μm in **A** and **D**, and 5 μm in **B** and **C**. In **A**, the numbers on the top are the number of midguts containing parasite / the number of midguts measured; IP: infection prevalence; red horizontal lines show the mean value of parasite numbers, and analyzed by Mann–Whitney test. All experiments in this figure were independently repeated three times, and the data are shown from one representative experiment.

### *Δatp7* ookinetes are eliminated within hours after midgut invasion

Following mosquito infection with blood containing gametocytes, ookinetes develop in the midgut lumen through an asynchronous process lasting 10 to 20 hours before midgut traversal. To allow synchronized ookinete invasion of epithelium, mosquitoes were infected by membrane feeding of ookinetes that were differentiated *in vitro* and purified using a Hemotek system [32]. Mosquitoes were fed for 20 minutes with a mixture of equal numbers of *Ogfp* and *OmCherry* ookinetes and midguts were dissected at 2.5, 4.5, and 8 h post feeding (pf) (Fig. 5A). At 2.5 h, comparable numbers of GFP^+^ ookinetes (*Ogfp*) and mCherry^+^ ookinetes (*OmCherry*) were detected, although the number of ookinetes among the midguts varied (Fig. 5B). Similar results were also obtained at 4.5 and 8 h pf (Fig. 5C and 6D). To delineate the precise timing of ookinete eradication, we disrupted the *atp7* gene in the *OmCherry* parasites (Fig. S3D) and performed co-feeding experiments with the resulting *OmCherry*/*Δatp7* mutant and the *Ogfp* ookinetes. Equal numbers of the GFP^+^ and mCherry^+^ ookinetes were observed at 2.5 h pf (Fig. 5E and F), indicating successful epithelium invasion by the ATP7-depleted ookinetes. However, at 4.5 h pf, the number of mCherry^+^ ookinetes was dramatically lower compared with that of GFP^+^ ookinetes (GFP^+^: 13.8±1.4; mCherry^+^: 0.8±0.2) (Fig. 5G). By 8 h pf, mCherry^+^ ookinetes completely disappeared (Fig. 5H). These results suggest that the ATP7-depleted ookinetes are eliminated within 2-4 hours after midgut invasion.

**Figure 5.**
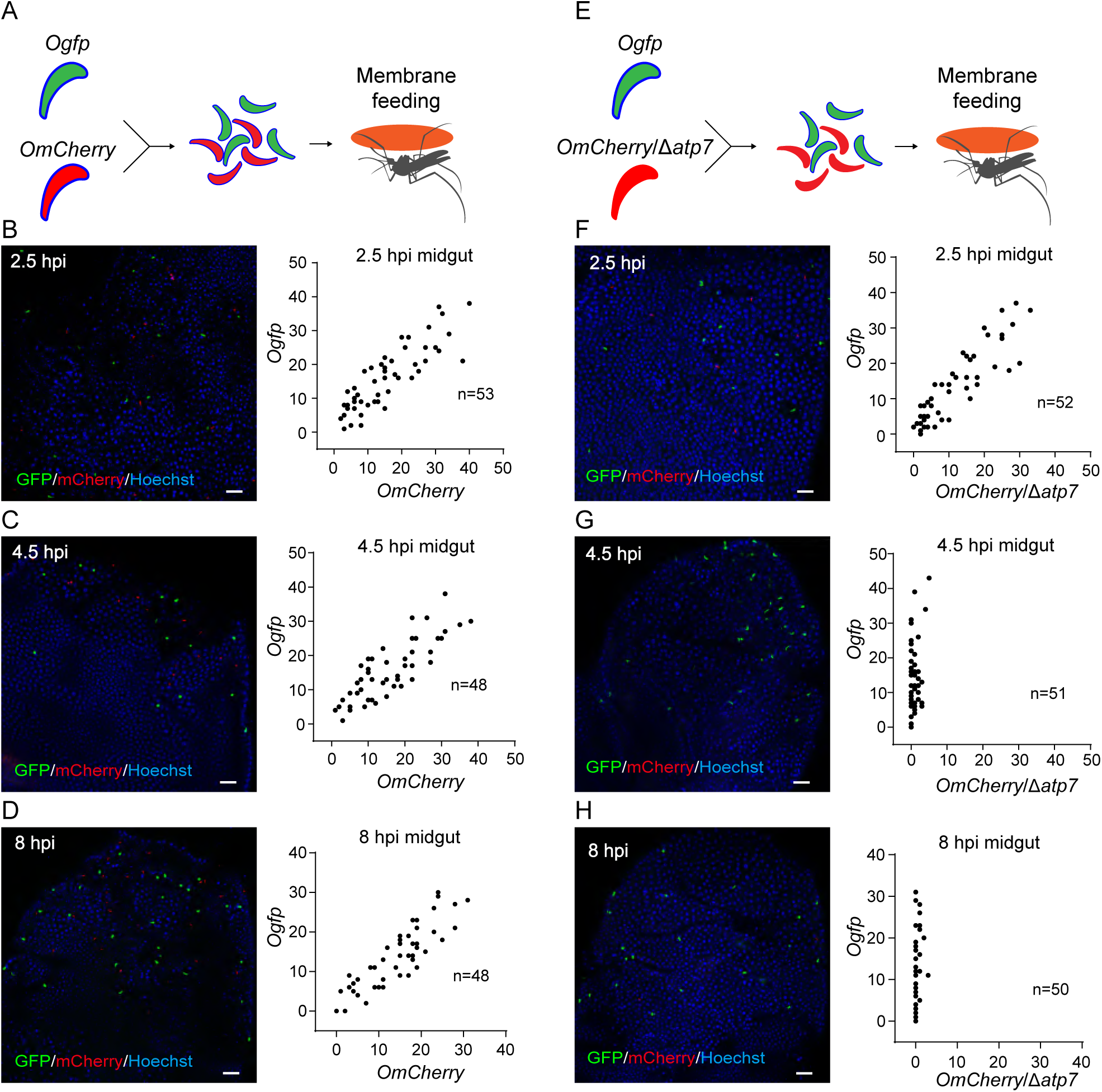
Δ*atp7* ookinetes are eliminated within hours after midgut invasion. **A**, **E**. Schematic of mosquito infection with *in vitro* purified ookinetes through membrane feeding. After co-feeding mosquitoes with equal number of ookinetes of *Ogfp* and *OmCherry* reporter strains (**A**), or *Ogfp* and *OmCherry*/Δ*atp7* strains (**E**), fluorescent ookinetes were counted in midguts dissected at 2.5, 4.5, and 8 h post membrane feeding (pf). **B-D**. *Ogfp* (green) and *OmCherry* (red) fluorescent ookinetes in co-infected midguts at 2.5 h (**B**), 4.5 h (**C**), and 8 h (**D**). The right panels display the quantification of GFP and mCherry fluorescent ookinetes. **F-H**. *Ogfp* (green) and *OmCherry*/Δ*atp7* (red) ookinetes in co-infected midguts at 2.5 h (**F**), 4.5 h (**G**), and 8 h (**H**). The right panels display the quantification of GFP and mCherry fluorescent ookinetes. n: number of midguts measured. Scale bar = 50 μm. Experiments in this figure were independently repeated twice, and the data are shown from one representative experiment.

**Figure 6.**
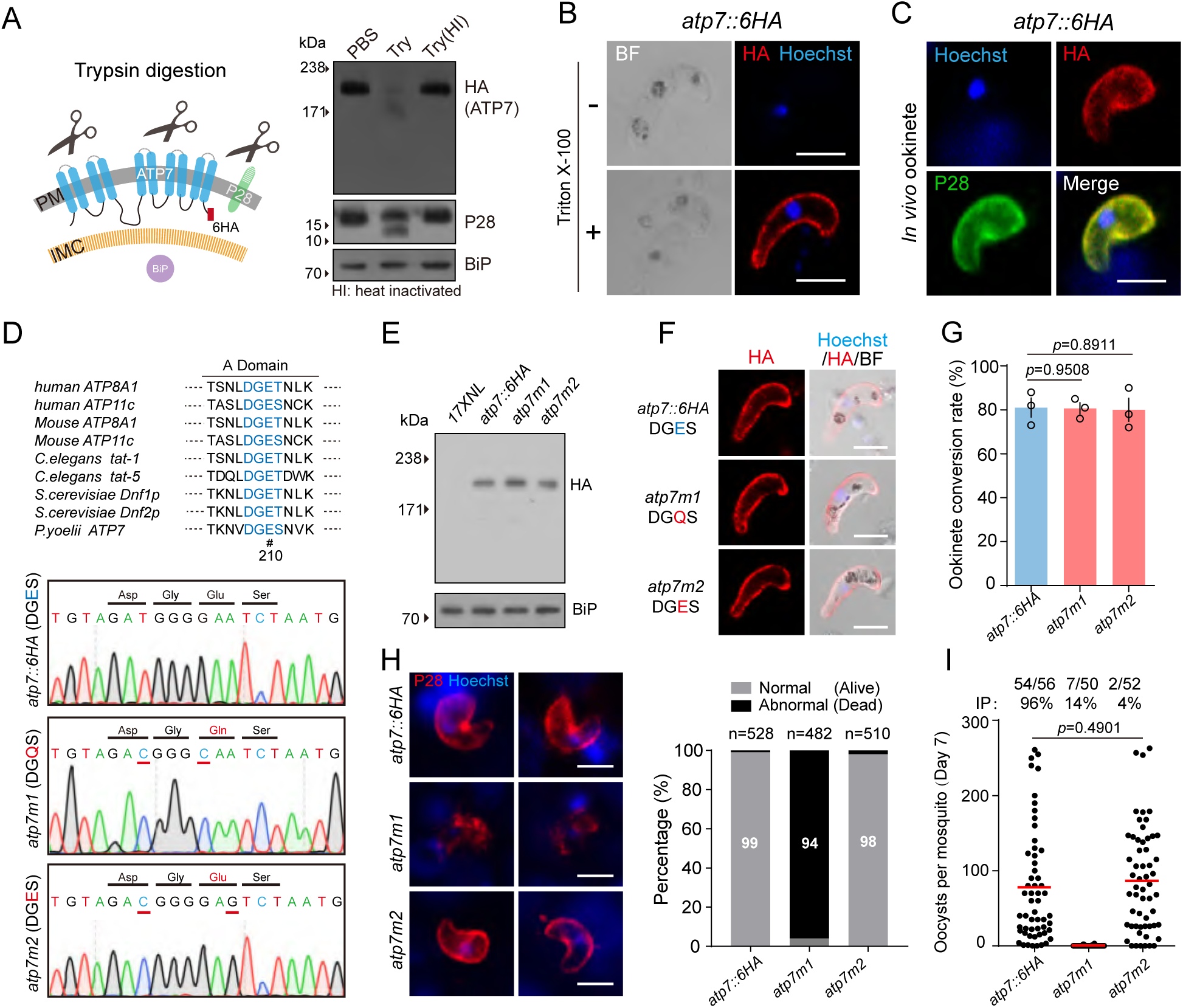
ATP7 is localized on ookinete plasma membrane and flippase activity is essential for its function. **A.** Immunoblot of ATP7, P28 (plasma membrane protein), and BiP (cytosolic protein) from *atp7::6HA* ookinetes treated with PBS, trypsin (Try), or heat-inactivated (HI) trypsin. Left panel is a schematic of plasma membrane (PM) and inner membrane complex (IMC) in ookinetes. Trypsin digests the surface proteins, but not intracellular proteins. **B.** IFA of ATP7 in the *atp7::6HA* ookinetes with or without cell permeabilization. **C.** ATP7 and P28 staining of *atp7::6HA* ookinetes from infected midguts. Midguts were permeabilized by Triton X-100. **D.** Generation of mutant parasites with impaired flippase activity of ATP7. Upper panel: conserved residues (blue letters) in actuator (A) domains critical for flippase activity of P4-ATPase. P4-ATPase proteins from *human*, *mouse*, *C. elegans*, *S. cerevsiae* and *P. yoelii* are shown. Residue E210 is marked with #. Bottom panels: DNA sequencing results confirming the E210Q substitution in *atp7m1* and E210E in *atp7m2* mutants. **E.** Immunoblot of ATP7 in ookinetes of *atp7::6HA*, *atp7m1*, and *atp7m2* parasites. BiP was used as a loading control. **F.** IFA of ATP7 in the ookinetes of the modified parasites. Ookinetes were permeabilized by Triton X-100. **G.** *In vitro* ookinete differentiation of the modified parasites. Data are shown as mean ± SEM of three independent experiments, two-tailed unpaired Student’s t test. **H.** Parasite plasma membrane and nucleus staining from infected midguts at 24 h pi. Right panel shows the percentage of normal (alive) and abnormal (dead) parasites. n is the number of parasites counted. **I.** Day-7 midgut oocysts in mosquitoes infected with the modified parasites. Numbers on the top are the numbers of midguts carrying oocysts / the number of midgut dissected; IP: infection prevalence; red horizontal lines show the mean parasite numbers. Scale bar = 5 μm in all images. Experiments in this figure were repeated three times, and the data are shown from one representative experiment.

### Elimination of *Δatp7* ookinetes is independent of mosquito complement system

We next asked how the *Δatp7* ookinetes get eliminated in the midgut. Mosquito complement-like immunity plays a major role in eliminating parasites during midgut traversal [3]. Is this immunity responsible for the elimination of *Δatp7* ookinetes? We infected *An. stephensi* mosquitoes in which the gene encoding a key component of the complement-like system, TEP1 [3], was silenced by injection with double stranded RNA (dsRNA) (Fig. S5A). Compared to the control GFP dsRNA, TEP1 silencing slightly increased (but not significantly) the number of WT oocysts. However, *Δatp7* oocyst formation was not restored in midguts neither at 2 nor 7 d pi after TEP1 silencing (Fig. S5B and C). Silencing of LRIM1 or APL1, two other critical complement components [8], also failed to reverse *Δatp7* ookinete elimination (Fig. S5D-G). These data suggest that the elimination of *Δatp7* ookinetes in midgut is independent of mosquito complement immunity.

To test whether mosquito hemocytes are involved in *Δatp7* ookinete elimination, we depleted phagocytic hemocytes in mosquito hemocoel using clodronate (CLD) liposomes [10] (Fig. S5H and I). CLD-treated mosquitoes were infected with blood containing *Ogfp* or *Ogfp*/*Δatp7* gametocytes. As expected [10], phagocyte depletion increased the number of *Ogfp* midgut parasites 24 h pi (Fig. S5J). However, it did not rescue the *Ogfp*/*Δatp7* defect (Fig. S5J). These results suggest that *Δatp7* ookinete elimination is not mosquito hemocyte-mediated.

### Ookinete microinjection into the mosquito hemocoel rescues the *Δatp7* defects

*Δatp7* ookinetes are completely eliminated during midgut traversal and they fail to further develop into oocysts and sporozoites. However, it is not clear whether ATP7 deficiency also affects the intrinsic developmental program of oocyst and sporozoite differentiation. By bypassing the physical barrier of the midgut, WT ookinetes injected into the hemocoel can develop into oocysts and salivary gland sporozoites in mosquitoes [33, 34]. At 24 and 48 h after injection of 690 cultured *Ogfp* and *Ogfp*/*Δatp7* ookinetes (Fig. S6A), comparable numbers of spherical GFP^+^ oocysts were detected in the hemocoel of the abdominal and thoracic cavities (Fig. S6B and C). Notably, the injected *Ogfp*/*Δatp7* ookinetes developed into approximately equal numbers of salivary gland sporozoites as the *Ogfp* ookinetes at 13 d post injection (Fig. S6D).

We also evaluated ookinete-to-oocyst transformation using an *in vitro* culture system [35]. As early as 5 h, GFP^+^ crescent-shaped *Ogfp* ookinetes protruded onto the outer convex edge, forming a snail-like stage described as a transforming ookinete [36], and further expanded to a spherical oocyst with a diameter of 7-8 μm at 24 h and 10 μm at 48 h (Fig. S6E). Depletion of ATP7 had no effect on this oocyst transformation process (Fig. S6F). Together, these experiments show that ATP7 is not required for differentiation into oocysts and sporozoites. Rather, it is required for protection of the ookinete from destruction in the midgut epithelial cells.

### ATP7 is localized in the ookinete plasma membrane and flippase activity is essential for its function

The ookinete pellicle harbors two membrane structures: the parasite plasma membrane (PPM) and beneath this the inner membrane complex (IMC). We sought to determine if ATP7 is localized to either of these membrane structures. We reasoned that if ATP7 is localized in the PPM, then trypsin digestion of the extracellular segment is capable of degrading ATP7. Indeed, immunoblotting with an HA antibody detected a band corresponding to the HA-tagged ATP7 (∼205 kD) in cell lysates prepared from *atp7::6HA* ookinetes treated with PBS- or heat-inactivated trypsin. Trypsin treatment completely abolished the immunoblot signal (Fig. 6A). It is worth noting that the PPM-residing protein P28 (but not the ER protein BiP) was efficiently cleaved, which validated our approach (Fig. 6A). In agreement with its predicted topology, the 6HA-tagged ATP7 could only be labeled if the ookinetes are permeabilized by Triton X-100 (Fig. 6B). ATP7 was also colocalized with P28 in the ookinetes (Fig. 6C). Together, these results indicate that ATP7 is localized in the ookinete plasma membrane.

To test whether a conserved flippase motif within ATP7 is required for its function, we generated parasites carrying *atp7* mutations that are predicted to compromise its flippase activity but not subcellular localization. It is known in other organisms that the conserved motif Asp-Gly-Glu-Ser (DGES) in the actuator domain are critical for the catalytic activity of P4-ATPases (Fig. 6D), and that mutations in these residues abolish the flippase activity [37]. Accordingly, we replaced E210 with Q in the *atp7::6HA* strain, in an attempt to generate an enzymatically dead mutant designated as *atp7m1* (Fig. 6D). A control strain (*atp7m2*) was also generated with a silent mutation still encoding “DGES” (Fig. 6D). The E210Q substitution had no effect on the protein level or localization of ATP7 in *atp7m1* ookinetes compared to the parental strain (Fig. 6E and F). Importantly, the *atp7m1* parasites produced comparable levels of ookinetes as the *atp7::6HA* strain (Fig. 6G), but were eradicated during midgut traversal (Fig. 6H) and therefore developed no oocysts in the mosquitoes (Fig. 6I). These results suggest that ATP7 is a functional flippase and that its activity is required for midgut traversal of ookinetes.

### ATP7 flips phosphatidylcholine

To gain further evidence that ATP7 is a functional flippase and to determine its preferred substrate, we compared the uptake of labelled phospholipids in WT and ATP7-depleted ookinetes. Using a phospholipid uptake assay established previously for use with mammalian P4-ATPases [38], we analyzed the flippase activity of ATP7 towards four phospholipids, including phosphatidylserine (PS), phosphatidylethanolamine (PE), phosphatidylcholine (PC), and sphingomyelin (SM), that were labeled with fluorescent NBD. Ookinetes were incubated in the presence of NBD-PS, PE, PC, or SM for 60 min at 22 °C, followed by removal of fluorescent unincorporated phospholipids and those attached to the ookinete surfaces. Uptake of PS, PE, PC, or SM occurred in the WT ookinetes in medium supplied with ATP, but did not occur in medium depleted of ATP (Fig. 7A and B). This is in agreement with the fact that P-type ATPases utilize energy from ATP hydrolysis to transport substrates across biological membranes [39]. Notably, only uptake of PC is impaired in the ATP7-depleted ookinetes compared to the WT ookinetes (Fig. 7C and D). Furthermore, we tested the uptake of PC in ookinetes of the *atp7m1* and *atp7m2* parasites. The enzymatic dead mutant *atp7m1* had impaired PC uptake, similar to that of *Δatp7* ookinetes, while the *atp7m2* ookinetes behaved normally (Fig. 7E and F), further confirming that ATP7 is a functional P4-ATPase and that it flips PC in ookinetes.

**Figure 7.**
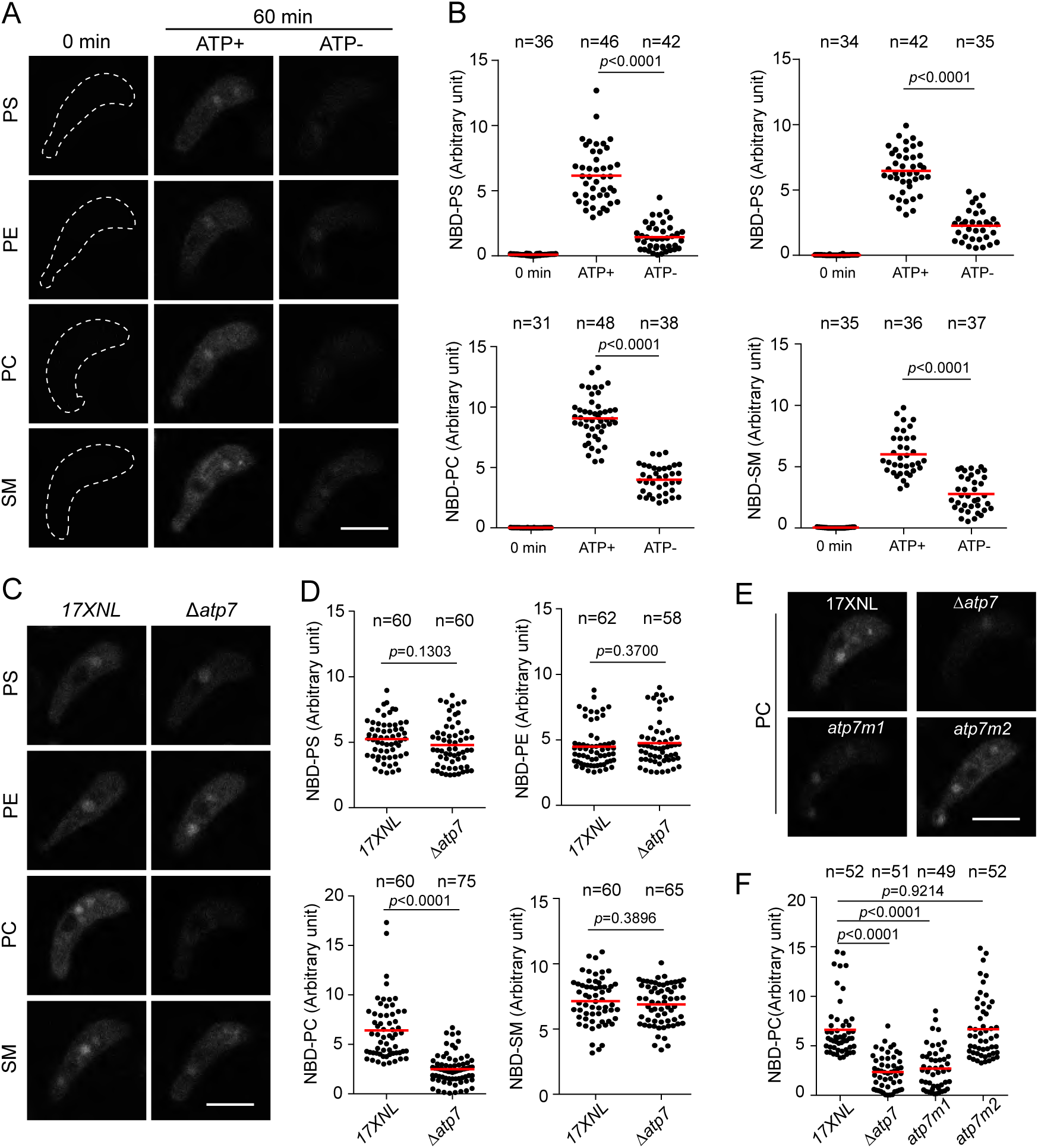
ATP7 flips phosphatidylcholine (PC). **A.** Phospholipid uptake by 17XNL ookinetes. Ookinetes were pre-incubated in HBSS-glucose medium (ATP+) or ATP-depleted medium (ATP-) for 60 min at 22 °C and then supplemented with different fluorescently-labeled NBD-phospholipids (PS, PE, PC, and SM) for 60 min at 22 °C. After washing with fatty acid-free BSA to remove unincorporated phospholipids and those attached to the ookinete surfaces, the ookinetes were imaged by confocal microscopy. Ookinetes with no supplementation of NBD-phospholipids (0 min) served as controls. **B.** Quantification of fluorescent signals of ookinetes in **A**. **C.** Phospholipid uptake by 17XNL and Δ*atp7* ookinetes. **D.** Quantification of fluorescent signals of ookinetes in **C**. **E.** NBD-PC uptake by 17XNL, Δ*atp7*, *atp7m1*, and *atp7m2* ookinetes. **F.** Quantification of fluorescent signals of ookinetes in **E**. Scale bar = 5 μm in all images. Red lines show mean values. n: the number of ookinetes analyzed in each group. Two-tailed unpaired Student’s t-test. Experiments in this figure were repeated three times, and the data are shown from one representative experiment.

It is not currently possible to explore whether PC is exposed at the PPM surface in the ATP7-depleted ookinetes because no commercial probe is available. However, staining living ookinetes with either Annexin V (PS probe) or Duramycin (PE probe) detected no fluorescent signal in either WT or *Δatp7* parasites (Fig. S7A and B), excluding surface exposure of PS and PE in ATP7-deficient ookinetes.

### ATP7 colocalizes and interacts with the CDC50C co-factor in ookinetes

P4-ATPase flippase function is reliant on its interaction with a CDC50 co-factor [40] (Fig. 8A). We investigated whether any of three CDC50 paralogues in malaria parasites interact with ATP7. In *P. yoelii* these are CDC50A: PY17X_0619700, CDC50B: PY17X_0916600, and CDC50C: PY17X_0514500 [23]. Of these, only CDC50C showed peripheral localization in ookinete [23]. CDC50C is predicted to contain two transmembrane helices and a large exocytosolic loop (Fig. 8B). Similarly as ATP7, CDC50C is sensitive to ookinete trypsin treatment and permeabilization (Fig. 8C and D), indicating PPM localization of CDC50C in ookinetes of the tagged parasite *cdc50c::6HA*. To determine whether CDC50C interacts with ATP7, we generated a double-tagged strain *atp7::6HA/50c::3V5* (Fig. S1), wherein ATP7 and CDC50C were tagged with 6HA and 3V5 respectively. ATP7 and CDC50C showed marked colocalization at the cell periphery in the mature ookinetes (Fig. 8E). This colocalization was also observed in another independent double-tagged strain *atp7::6HA/50c::4Myc* (Fig. 8F). Furthermore, immunoprecipitation using an HA antibody indicated that ATP7 is bound to CDC50C in the *atp7::6HA/50c::4Myc* ookinete lysates (Fig. 8G). Lastly, we performed a proximity ligation assay (PLA), an immunohistochemical method to detect proteins interaction *in situ* [41]. A robust PLA signal was detected at the ookinete periphery in the *atp7::6HA/50c::4Myc* parasites when both anti-HA and anti-Myc primary antibodies were present (Fig. 8H), indicative of ATP7 and CDC50C interaction. Noteworthy, no signal was observed in either *atp7::6HA* or *atp7::6HA/50c::3V5* ookinetes. These results demonstrate that ATP7 co-localizes and interacts with CDC50C in the PPM of ookinetes.

**Figure 8.**
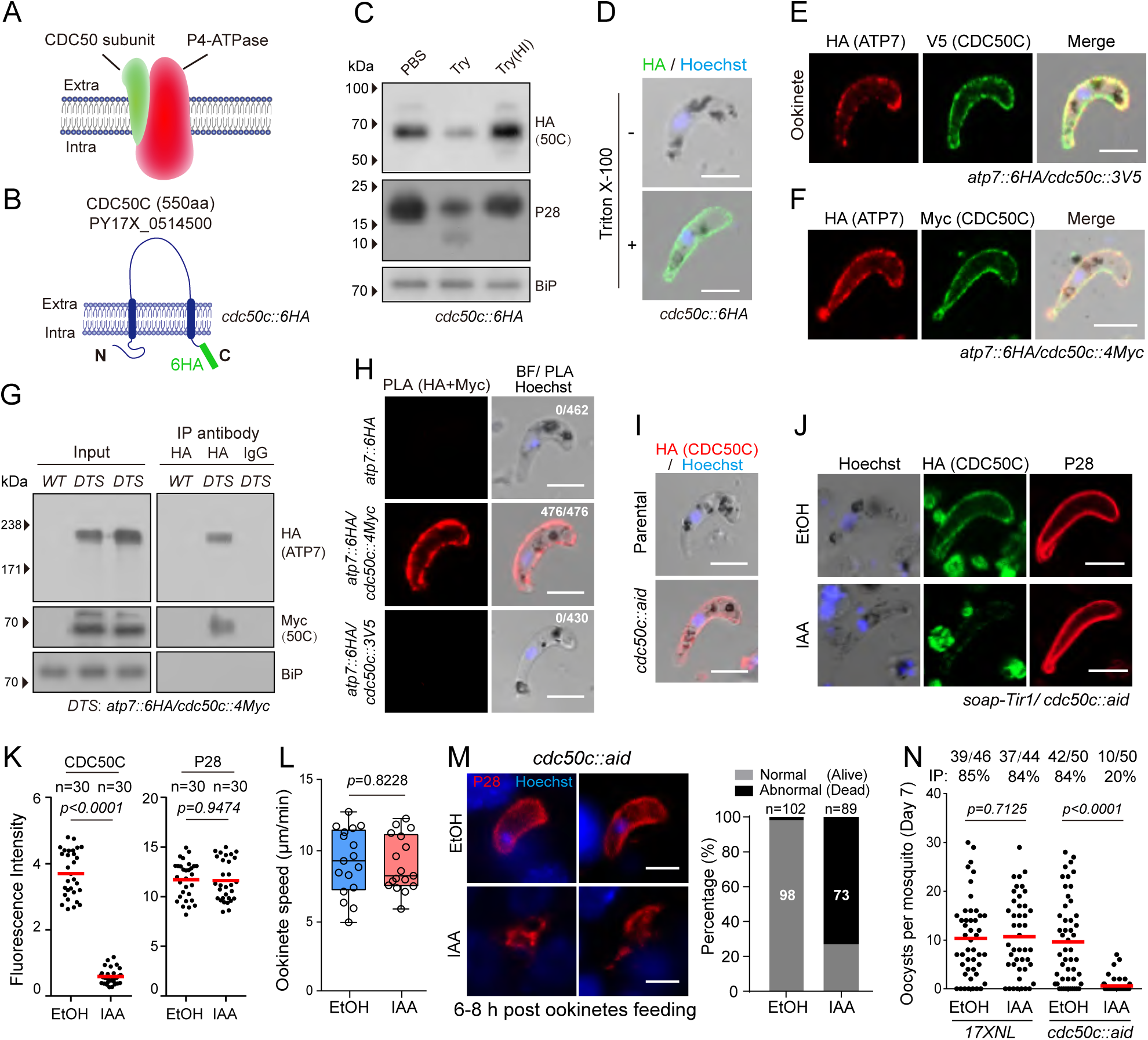
CDC50C is a ATP7 co-factor for flippase function. **A.** Diagram of the eukaryotic flippase complex made of a P4-ATPase and a CDC50. **B.** Topology of the *P. yoelii* CDC50C protein (PY17X_0514500) in ookinete plasma membrane of the tagged parasite *cdc50c::6HA*. **C.** Immunoblot of CDC50C, P28, and BiP of *cdc50c::6HA* ookinetes treated with PBS, trypsin (Try), or heat-inactivated (HI) trypsin. **D.** IFA of ATP7 in *atp7::6HA* ookinetes with or without Triton X-100 permeabilization. **E**-**F**. Two-colored IFA of ATP7 and CDC50C proteins in ookinetes of two double-tagged parasite strains: *atp7::6HA*/*cdc50c::3V5* (**E**) and *atp7::6HA*/*cdc50c::4Myc* (**F**). **G.** Co-immunoprecipitation assay of ATP7 and CDC50C proteins from ookinetes of the double tagged strain *DTS* (*atp7::6HA*/*cdc50c::4Myc* strain). **H.** Proximity Ligation Assay (PLA) detecting protein interaction between ATP7 and CDC50C in *atp7::6HA*/*cdc50c::4Myc* ookinetes. On the right panels: number of cell displaying signal / the number of cell counted. **I.** Generation of the *cdc50c::aid* strain with endogenous CDC50C tagged with an AID::6HA motif for auxin-induced protein degradation. The parental is an engineered strain *soap*-*Tir1* stably expressing the *Tir1* in the ookinetes. **J.** IAA-induced depletion of CDC50C protein in the ookinetes of *cdc50c::aid* strain. Ethanol treatment as control. **K.** Quantification of fluorescent signal intensity of proteins (CDC50C and P28) in **J**. Red lines show the mean value. n is the number of ookinetes analyzed in each group. Two-tailed unpaired Student’s t-test. **L.** Ookinete gliding motility *in vitro*. **M.** Parasite plasma membrane and nucleus staining in midguts infected with *in vitro* purified and IAA-treated ookinetes through membrane feeding. Midguts were dissected at 6-8 h post feeding. The right panel shows the percentage of normal (alive) and abnormal (dead) parasites. n is the number of parasites counted. **N.** Day-7 midgut oocyst numbers in mosquitoes infected with the modified parasites. Numbers on the top are the number of midguts carrying oocyst / the number of midguts dissected; IP: infection prevalence; red horizontal lines show the mean value of parasite numbers. Scale bar = 5 μm in all images. Experiments in **B**, **C**, **D**, **E**, **F**, **H**, **I**, **J**, and **L** were repeated three times, and experiments in **M** and **N** were repeated twice. The data are shown from one representative experiment.

### CDC50C depletion in ookinetes phenocopies ATP7 deficiency during mosquito transmission

To investigate the potential function(s) of CDC50C in mosquito transmission, we attempted to disrupt the *cdc50c* gene but failed to obtain mutants, suggesting an essential role of CDC50C in parasite asexual blood stages. This observation is in agreement with the recent global knockout screening results in *P. falciparum* and *P. berghei* [42, 43]. Therefore, we applied an auxin-inducible degron (AID)-based protein degradation system we recently adapted in *P. yoelii* (manuscript in revision), which allows rapid depletion of a target protein fused to an AID motif with the aid of auxin/IAA [44]. The C-terminus of the endogenous *cdc50c* locus was tagged with the sequence encoding the AID::6HA in the *soap*-*Tir1* strain (manuscript in revision) (Fig. S1). The resulting *cdc50c::aid* parasite exhibited the expected cell periphery localization of CDC50C in ookinetes (Fig. 8I), indicating no marked effect of AID tagging on CDC50C localization. IAA treatment (1 mM for 1 h) dramatically reduced the CDC50C protein abundance, but had little impact on P28 levels (Fig. 8J-K). Of note, *in vitro* gliding motility of the IAA-treated ookinetes is comparable to controls (Fig. 8L). Furthermore, we employed a membrane feeding experiment using ookinetes treated with either IAA or EtOH to test the effect of CDC50C depletion on parasite transmission. Notably, IAA-treated ookinetes were lysed during midgut epithelium traversal (Fig. 8M). Consistent with these observations, IAA-treated ookinetes developed a much lower number of oocysts in mosquito midguts compared to the controls at 7 day pf (Fig. 8N). Together, these results strongly suggest a critical role of CDC50C in safeguarding ookinetes during midgut traversal, similar to that of its binding partner ATP7.

## Discussion

In this study, we identified a *Plasmodium* flippase complex comprising a catalytic P4-ATPase ATP7 (α-subunit) and a CDC50C (β-subunit) co-factor in the rodent malaria parasite *P. yoelii*. ATP7 and CDC50C are localized in the plasma membrane of mature ookinetes, facilitating ookinete survival during its traversal of the mosquito midgut epithelium. Whilst ATP7- or CDC50C-depleted ookinetes are motile and capable of midgut invasion, they are quickly eliminated by the midgut epithelium, leading to failure of parasite transmission. It is noted that successful gene disruption of *atp7* orthologs have been reported in genome-wide gene knockout screens of *P. falciparum* and *P. berghei* [42, 43]. However, an attempt to disrupt *atp7* ortholog in another gene knockout screen of *P. berghei* was not successful using conventional homologous recombination method [45], possibly resulting from variable rate of recombination at this locus among parasite specices and strains tested.

As a putative flippase based on protein sequence homology, ATP7 was predicted to transport phospholipids from the outer leaflet to the inner leaflet of the lipid bilayer but the identity of the lipid substrate was unknown. Our experiments revealed that ATP7 is a functional P4-ATPase essential for midgut traversal of malaria parasite. Furthermore, in the phospholipid uptake assay, WT ookinetes were able to take all four phospholipids tested (PS, PE, PC, and SM) from the medium into the ookinetes. However, the uptake of only PC is impaired in the ATP7-depleted ookinetes, suggesting that PC is the specific substrate for ATP7. Consistent with these data, the flipping of PC by ATP7 was abrogated by a point mutation in its enzymatic active site. Therefore, we speculate that in the absence of APT7, PC is exposed at the outer plasma membrane of the ookinete (graphic abstract in Fig S8). Direct demonstration of PC exposure in the *Δatp7* ookinetes is not currently possible for lack of a PC-specific chemical probe.

How does a defect in ATP7/CDC50C-mediated PC translocation across plasma membrane results in ookinete elimination during midgut traversal? P4-type ATPases are involved in the maintenance of phospholipid asymmetry which might directly or indirectly protect the parasites from the environmental stress imposed by the hosts. Conceivably, an altered distribution of phospholipids may compromise the structure and permeability of the plasma membrane and therefore cell viability. Surprisingly, our data show that WT and the ATP7-depleted ookinetes exhibit similar cell morphology and survival under different stress conditions (Fig. S2D-I). Moreover, *Δatp7* ookinetes are also indistinguishable from WT regarding microneme secreation, cell pellicle and apical structures, and gliding motility. Instead, once the *Δatp7* ookinetes invade the midgut epithelium, the PC exposed on the ookinetes surface possibly functions as a pathogen-associated molecular pattern (PAMP), which is recognized by the pattern recognition receptors (PRRs) of the mosquito host [46], inducing rapid ookinete elimination (graphic abstract in Fig S8). Notably, surface display of PS due to the absense of flippase activity marks apoptotic cells for consumption by phagocytes[47]. However, the exact mechanisms underlying ATP7-deficient ookinete-induced immunity within the mosquito midgut epithelium requires further investigation.

During the *Plasmodium* life cycle, three invasion events need to occur: sporozoite invasion of hepatocytes and merozoite invasion of erythrocytes in the verterbrate host, and ookinete invasion of the midgut in the mosquito. All three invasive forms share cellular machinery that drives these processes [48]. Remarkably, merozoite and sporozoite invasion are accompanied by invagination of the host cell plasma membrane, resulting in the formation of a parasitophorous vacuole (PV). The parasites undergo asexual proliferation inside the PV, where the PV membrane (PVM) functions as a barrier between the parasite and the host cell cytoplasm and protects the parasite from anti-*Plasmodium* responses by the host cells [49, 50]. However, no PV formation occurs during ookinete invasion of the midgut epithelium [51]. This scenario of direct contact between the invading ookinete and epithelium cytoplasm raises the possibility that ookinetes could be recognized and targeted by the mosquito intracellular immunity. Consistent with this hypothesis, co-infection experiments showed that ATP7 on the surface of WT ookinetes does not confer protection to the co-invading ATP7-deficient ookinetes. Therefore, protection of the ookinete by ATP7 is not due to a systemic mechanism in the midgut but an intrinsic mechanism to the individual ookinetes whithin the epithelium.

It is well established that the ookinete surface protein P47 is a key molecule that protects the *Plasmodium* ookinetes from mosquito complement-dependent immune attack during the midgut traversal [5]. P47 inhibits mosquito Janus kinase signaling as the ookinetes come in close contact with the mosquito hemolymph [17]. However, in our case, genetic and chemical ablation of either mosquito complement or hemocyte cellular immunity failed to rescue the aberrant eradication of *Δatp7* ookinetes. Therefore, *Plasmodium* ATP7/CDC50C-mediated protection of ookinetes likely represents a new mechanism of malaria parasite evasion from the mosquito host immune system.

## Acknowledgments

We thank Dr. Bo Wang, Dr. David Baker, and Dr. Marcelo Jacobs-Lorena for the comments on this manuscript. This work was supported by the National Natural Science Foundation of China (31772443, 31872214, 31970387), the Natural Science Foundation of Fujian Province (2019J05010), and the 111 Project sponsored by the State Bureau of Foreign Experts and Ministry of Education of China (BP2018017).

## Author contributions

Y.ZK., S.Y., and G.H. generated the modified parasites, conducted the phenotype analysis, IFA assay, image analysis, mosquito experiments, and biochemical experiments, Y.SZ. conducted the mosquito experiments, Y.J. and C.HT, supervised the work. Y.ZK, C.HT, and Y.J. analyzed the data, and Y.J. wrote the manuscript.

## Material and Method

### Mice and mosquitoes usage and ethics statement

All animal experiments were performed in accordance with approved protocols (XMULAC20140004) by the Committee for Care and Use of Laboratory Animals of Xiamen University. The ICR mice (female, 5 to 6 weeks old) were purchased from the Animal Care Center of Xiamen University and used for parasite propagation, drug selection, parasite cloning, and mosquito infection. The mosquito of *An. stephensi* (Hor strain) was reared at 28°C, 80% relative humidity and at a 12h light/dark cycle in the standard insect facility. Adult mosquitoes were maintained on a 10% sucrose solution.

### Plasmid construction and parasite transfection

CRISPR/Cas9 plasmid pYCm was used for all the parasite genetic modification. To construct vectors for gene editing, we amplified 5’ and 3’ genomic sequence (400 to 500 bp) of target genes as homologous arms using specific primers (Table S2) and inserted the sequences into specific restriction sites in pYCm. Oligonucleotides for guide RNAs (sgRNAs) (Table S2) were mixed in pairs, denatured at 95°C for 3 min, annealed at room temperature for 5min, and ligated into pYCm. The sgRNAs were designed to target the coding region of a gene (Table S2) using the online program EuPaGDT. DNA fragments encoding 6HA, 4Myc, 3V5, and GFP or mCherry were inserted between the left and right arms in frame with gene of interest. For each gene, two sgRNAs were designed to target sites close to the C- or N-terminal part of the coding region. Infected red blood cells (iRBC) were electroporated with 5∼10 μg plasmid DNA using Lonza Nucleofector described previously [21]. Transfected parasites were immediately intravenously injected into a naïve mouse and were exposed to pyrimethamine (6 mg/ml) 24h post-transfection.

### Genotypic analysis of genetic modified parasites

All transgenic parasites were generated from *P. yoelii* 17XNL strain and are listed in Table S1. Blood samples from infected mice were collected from the orbital sinus, and blood cells were lysed using 1% saponin in PBS. Parasite genomic DNAs were isolated from blood stage parasites using DNeasy blood kits (#69504, QIAGEN). For each parasite, both 5’ and 3’ homologous recombination events were detected using specific PCR primers (see Fig. S1). The PCR results for parasite transfection, selection, cloning and verification of genetic modifications are summarized in Fig. S1. PCR products from some modified parasites were DNA sequenced. All the primers used in this study are listed in Table S2. Parasite clones with targeted modifications were obtained after limiting dilution. At least two clones for each gene-modified parasite were used for phenotype analysis.

### Negative selection with 5-fluorouracil

Parasites subjected to sequential modification were negatively selected with 5-Fluorouracil (5FC, Sigma, F6627) to remove episomal plasmid. 5FC (2 mg/ml) in drinking water in a dark bottle was provided to mice for 8 days with a replacement on day 4. Clearance of episomal plasmid in parasites after negative selection was confirmed by checking the parasite survival after reapplying pyrimethamine pressure (6 μg/ml) in new infected mice.

### Parasite proliferation assay in asexual mouse blood stage

1.0×10^5^ parasites were injected into tail vein of each of five naive ICR mice in each group. Parasitemia is monitored every two days through Giemsa staining of the thin mouse blood smears from day 2 to day 16 post infection. The parasitemia is calculated as the ratio of parasitized erythrocytes over total erythrocytes.

### Gametocyte induction in mouse

ICR mice were treated with phenylhydrazine (80 μg/g mouse body weight) through intraperitoneal injection. Three days post injection, each mouse was infected with 4.0 ×10^6^ parasites through intravenous injection. Gametocytemia usually peaks at day 3 post infection. Male and female gametocytes were counted after Giemsa staining of blood smears. Male and female gametocytemia were calculated as the ratio of male and female gametocytes over the parasitized erythrocytes. These experiments were repeated three times independently.

### Gametocyte activation to gamete *in vitro*

Two and a half microliters of mouse tail blood with 4-6% gametocytemia were added to 100 μl exflagellation medium (RPMI 1640, 10% fetal calf serum/FCS, 100 μM xanthurenic acid/XA, and pH 7.4) containing 1 μl of 200 unit/ml heparin for 10 min at 22°C. For male gametocyte activation, the exflagellation centers (EC) and total RBC were counted in a hemocytometer under light microscope. The percentage of RBCs containing male gametocytes was calculated from Giemsa-stained smears, and the number of ECs per 100 male gametocytes was calculated as male gamete formation rate. For female gametocyte activation, P28 positive female gametocytes after staining with P28-antiserum and RBC were counted. The percentage of RBCs containing female gametocytes was calculated from Giemsa-stained smears, and the number of P28 positive female gametocytes per 100 female gametocytes was calculated as female gamete formation rate.

### *In vitro* ookinete differentiation

*In vitro* ookinete differentiation was performed as described previously [23]. Briefly, 1 ml of mouse blood with 4-6% gametocytemia were collected via orbital sinus and immediately transferred to a 10 cm cell culture dish (Corning, cat#801002) containing ookinete culture medium (RPMI 1640, 10% FCS, 100 μM XA, 25mM HEPES, and pH 8.0). The cultures were incubated at 22°C for 12-14 hr to allow gametogenesis, gamete fertilization, and ookinete differentiation. Ookinete formation was monitored by Giemsa-staining of culture smears. Ookinete conversion rate was calculated as the number of ookinetes per 100 female gametocytes. For time-course analysis of ookinete differentiation, parasites from 2, 4, 6, 8, 12 hr culture were collected. Ookinetes were purified using ACK lysing method [23]. Briefly, the cultured ookinetes were collected by centrifugation and transferred into ACK lysing buffer (A1049201, ThermoFisher Scientific) on ice for 8 min. After erythrocytes lysis, the remaining ookinetes were isolated via centrifugation and washed twice with PBS. The ookinetes were examined and counted on the hemocytometer under 40× objective lens. Samples containing > 80% ookinete population were used for further analysis.

### *In vitro* ookinete motility assay

Ookinete gliding motility was tested as previously described [23]. All procedures were performed in a temperature-controlled room with 22°C. Briefly, 20 μl of mature ookinete cultures were mixed with 20 μl of Matrigel (#356234, BD) on ice. The mixtures were transferred onto a slide, covered with a coverslip, and sealed with nail varnish. The slide was placed at 22°C for 30 min before observation under microscope. After tracking a gliding ookinete under view of 40× objective lens, time-lapse videos (1 frame/20 sec) were taken to record ookinete movement for 20 min using a Nikon ECLIPSEE100 microscope fitted with an ISH500 digital camera and ISCapture v3.6.9.3N software (Tucsen). Time-lapse movies were analyzed with Fiji software and the Manual Tracking plugin. Motility speed was calculated by dividing the gliding distance of ookinete by the time. The experiments were repeated three times independently with 15-20 ookinetes tested for each parasite in each time.

### Mosquito feeding and transmission assay

Thirty female mosquitoes were allowed to feed on an anesthetized mouse carrying 8-10% gametocytemia for 30 min at 22°C. Mosquito midguts were dissected at the indicated time (1, 2, 3, 5, or 7 day) post blood-feeding and stained with 0.1% mercurochrome for visualizing the midgut oocysts. Mosquito salivary glands were dissected on day 14 post blood-feeding, and the number of sporozoites per mosquito was calculated. For parasite transmission from mosquito to mouse, twenty infected mosquitoes were allowed to feed on a naïve mouse and the parasites emerging in mouse blood were examined.

### Ookinete microneme secretion

Ookinete microneme secretion was analyzed as previously reported [44]. Briefly, about 2.0×10^6^ of ookinetes were purified from *in vitro* culture using the LS magnetic column (#130-042-401, Miltenyi Biotec) and incubated in 500 ul of PBS for 4 hr at 22°C. The cells were spun down at 750×g and the supernatant collected, filtered through a filter (0.45 µm, SLHP033RS, Millipore) and subject for western blot assay.

### Parasite genetic cross

The phenylhydrazine pre-treated mice were co-infected with same amount (2.0×10^6^) of each parasite for genetic crosses (Δ*nek4*×Δ*map2*, Δ*atp7*×Δ*map2* and Δ*atp7*×Δ*nek4*). On day 3 post infection, fifty female mosquitoes were allowed to feed on gametocyte-containing mice for 30 min. Mosquito midguts were dissected on day 7 post blood-feeding and stained with 0.1% mercurochrome for oocyst counting.

### Mosquito membrane feeding with ookinetes

Purified ookinetes (2.0×10^6^) from *in vitro* culture were suspended in 1 ml of naive mouse blood. The blood mix was added to membrane feeder and fed to about sixty female mosquitoes for 30 min using the Hemotek (6W1, Hemotek Limited, England). After the feed, fully engorged mosquitoes were transferred to the new container and maintained under standard conditions. The mosquito midguts were dissected and observed at indicated time (2.5, 4.5, and 6 h post feeding). These experiments were repeated three times. To test the mosquito transmission of ookinetes with chemical depletion of CDC50C, the ookinetes treated with IAA or EtOH were immediately added to membrane feeder and fed to the mosquitoes.

### *In vitro* culture for oocyst transformation

The procedure and culture conditions were followed as previous study with some modifications [36]. The *in vitro* differentiated ookinetes were seeded into the 8-well LabTek chamber slide or the 48-well plate at a density of 1×10^4^ ookinetes per well and cultured in oocyst culture medium (85% Schneider’s medium, 15% FBS, 23.8 mM sodium bicarbonate, 44 mM PABA, 3.68 mM Hypoxanthine, and pH 8.0). Parasites were maintained at 22°C for 2 days, and the oocysts were monitored using the fluorescent microscope. These experiments were repeated three times with six replicate cell for each parasite in each test.

### Ookinete microinjection into mosquito

After ookinete purification from *in vitro* culture described above, each of forty mosquitoes (4 to 5 day old) was injected intra-thoracically with a 138 nl of PBS containing 690 ookinetes using a Nanoject II injector (Drummond Scientific, USA). The hemocoel oocysts within the mosquitoes were observed under the stereoscopic fluorescence microscope (M165FC, Leica) at indicated times post injection.

### Quantitative real-time PCR

Thirty mosquito midguts are pooled in each group per condition per experiment. Total RNA was isolated from homogenized midgut using the RNeasy Mini Kit (74106, Qiagen) and reverse-transcribed to complementary DNA (cDNA) using the Kit (KR116-02, TIANGEN). Transcript expression of target mosquito genes was quantified using SYBR Green Supermix (#1708882, Bio-RAD) and Bio-Rad iCycler iQ system (Bio-Rad, USA). The cycling conditions were as follow: 95°C for 20 sec followed by 40 cycles of 95°C for 3 sec; 72°C for 30 sec. The samples were run in triplicate and normalized to the *An. Stephensi rps7* gene (ASTE004816) using a ΔΔ cycle threshold-based algorithm. Arbitrary unit is provided to represent relative expression levels. PCR primers used are listed in Table S2.

### Double-stranded RNA synthesis and mosquito gene silencing

Double-stranded RNA (dsRNA) were synthesized by *in vitro* transcription using the MEGAscript RNAi Kit (AM1626, Ambion). The DNA templates for transcription were PCR-amplified from the mosquito cDNA using the primer pairs containing sequence of T7 polymerase promoter at the 5’-end (Table S2). For gene silencing, each of fifty mosquitoes (4 to 6 day old) was injected intra-thoracically with a 69 nl of dsRNA solution (3 μg/μl) using a Nanoject II injector. Ten mosquitoes were collected on 2 or 3 day post injection and the gene knockdown efficiency was evaluated using the real-time quantitative PCR before parasite infection. The experiments were repeated three times.

### Mosquito phagocyte depletion using clodronate liposome

Mosquito phagocyte depletion was performed according to a method recently developed [10]. Each naive female mosquito was injected intra-thoracically with 69 nl of control liposome (LP) or clodronate liposome (CLD) (#8901, Encapsula NanoSciences) using the Nanoject II injector. 100% LP and different concentrations (100%, 50%, and 20%) of CLD diluted with 1xPBS were initially tested to determine the efficacy on phagocyte depletion. The CLD (20%) was used in the subsequent experiments. Injection was performed on naïve mosquitoes one day prior to the infected-mouse feeding.

### Mosquito hemolymph perfusion and hemocyte counting

Mosquito hemolymph was collected using the perfusion method [10]. Two day post injection with liposome, the mosquito was injected intra-thoracically with a 10 µl of the anticoagulant solution (60% Schneider’s insect medium, 10% FCS, 30% citrate buffer, 98 mM NaOH, 186 mM NaCl, 1.7 mM EDTA, 41 mM citric acid, and pH 4.5) and was perfused through an incision made in the lateral abdomen. About 10 µl of hemolymph was collected from one mosquito.

### Mosquito hemocyte staining using CM-DiI

Hemocytes were visualized by staining with Vybrant CM-DiI (V22888, Life Technologies), a lipophilic dye specifically labeling mosquito hemocytes [10]. Mosquitoes pre-treated with liposome were infected with parasites. 48 h post infection, the mosquitoes were injected with 138 nl of 100 μM CM-DiI and incubated for 20 min at 22°C. Perfused hemolymph (10 μl) was collected onto a cover glass (#801010, NEST, China) and hemocytes were allowed to adhere to the slide for 30 min. Without washing, 4% paraformaldehyde was added to each well for fixation. The cells were incubated at room temperature for 30 min, washed three times with PBS, and observed under the fluorescence microscope (DM4B, Leica, Germany). Hemocytes per mosquito were quantified after counting 15 fields (40×) of microscope.

### Antibodies and antiserum

The primary antibodies used in this study include: rabbit anti-HA (3724S, CST, Western blot, 1:1000 dilution; IFA, 1:500 dilution), rabbit anti-Myc (2276S, CST, Western blot, 1:1000; IFA, 1:500) from, mouse anti-V5 (A01724-100, Genscript, Western blot, 1:1000, IFA, 1:500), mouse anti-HA (sc-57592, SantaCruz, IFA, 1:200), mouse anti-Myc (sc-40, SantaCruz, IFA, 1:200), mouse anti-α-Tubulin II (T6199, Sigma-Aldrich IFA, 1:1000); mouse anti-nitrotyrisine from (SAB5200009, Sigma, IFA, 1:1000); Biotin-LC-Duramycin (25690-100, Polysciences, IFA, 1:1000). The secondary antibodies include: goat anti-rabbit IgG HRP-conjugated and goat anti-mouse IgG HRP-conjugated secondary antibodies (Abcam, 1:5000); the Alexa 555 labeled goat anti-rabbit IgG, Alexa 555 labeled goat anti-mouse IgG, and Alexa 488 labeled goat anti-mouse IgG secondary antibodies from Thermo Fisher Scientific (1:500); Streptavidin-488 (Biosciences, IFA, 1:500). Antiserum of rabbit anti-P28 (Western blot, 1:1000; IFA, 1:1000), rabbit anti-GAP45 (IFA, 1:1000), and rabbit anti-BiP (Western blot, 1:1000) were prepared previously [23]. Other polyclonal antiserum were prepared by immunization of rabbit with synthetic peptides or recombinant protein as follow: *α*-enolase (KTYDLDFKTPNNDK, 1:1000), *α*-chitinase (HTEKQYKSLSHVDALC, 1:1000), *α*-CTRP (LNGGETPHNSNMEFENVENNDGIIEEENEDFEVIDANDPMW, 1:1000), *α*-WARP (CNKNNPSSLTSERKTTIKN, 1:1000), and *α*-SRPN6 (RRDQWRRQTCPHDED, 1:1000).

### Parasite immunofluorescence assay

Parasites of different stages were fixed using freshly prepared 4% paraformaldehyde (P6148, Sigma-Aldrich) in PBS for 15 min at room temperature and transferred to a 24-well cell plate containing a Poly-L-Lysine (#E607015, Sangon Biotech) pre-treated coverslip at the bottom. The fixed cells were then immobilized on the coverslip via centrifuging the plate at 2000 rpm for 10 min and washed twice with PBS. The fixed cells were permeabilized with 0.1% Triton X-100 PBS solution for 10 min at room temperature, washed with PBS three times, blocked in 5% BSA solution for 60 min at room temperature, and incubated with the primary antibodies diluted in 5% BSA-PBS at 4°C for 12 h. The coverslip was incubated with fluorescent conjugated secondary antibodies for 1 h at room temperature and washed with PBS three times. Cells were stained with Hoechst33342 (#62249, Thermo Fisher), mounted in 90% glycerol solution, and sealed with nail polish. All images were captured and processed using identical settings on a Zeiss LSM 780 confocal microscopy.

### Mosquito midgut immunofluorescence assay

Mosquitoes fed on parasite-infected mice were dissected and the blood bolus contents were removed. Midguts were fixed with 4% paraformaldehyde for 1 h in PBS at room temperature, permeabilized with 100% methanol for 20 min at -20°C, then blocked in 5% BSA solution for 1 h at room temperature. Next, midguts were incubated overnight with the primary antibodies or antisera (1:1000 dilution in 5% BSA), then with the Alexa555 or 488-conjugated secondary antibodies (1:1000 diluted in 5% BSA). Midguts were then stained with Hoechst33342, mounted in 90% glycerol solution, and sealed with nail polish. All images were captured and processed using identical settings on a Zeiss LSM 880 confocal microscope. To visualize the midgut in a Z-stack manner, midguts were fixed in 4% paraformaldehyde in PBS for 1 h at room temperature and stained with anti-P28 antiserum. Then the midguts incubated with 6.6 μM Alexa Fluor 488-conjugated phalloidin (#21833, Invitrogen) for 20 min at room temperature. Z-stacks were approximately 20 μm length to encompass the interface of epithelial layer and the basal lamina, and images were taken at intervals of 1 μm. The final images were obtained and analyzed using Zeiss ZEN 3.0 software.

### Protein extraction and immunoblot

Proteins were extracted from asexual blood parasites, gametocytes, and ookinete using buffer A (0.1% SDS, 1 mM DTT, 50 mM NaCl, 20 mM Tris-HCl, pH 8.0) and from mosquitoes using mosquito protein extraction buffer (8M urea, 2% SDS, 5% β-mercaptoethanol, 125Mm Tris-HCl, pH8.0) containing protease inhibitor cocktail and PMSF. After ultrasonication, the protein solution was kept on ice for 15 min before centrifugation at 14,000 g for 10 min at 4°C. The supernatant was lysed in Laemmli sample buffer stored at 4°C or immunoblotting. The protein samples were separated in SDS-PAGE and transferred to PVDF membrane that was blocked in TBST buffer with 5% skim milk and then incubated with primary antibodies. After incubation, the membrane was washed three times with TBST and incubated with HRP-conjugated secondary antibodies. The membrane was washed four times in TBST before enhanced chemiluminescence detection.

### Proximity ligation assay (PLA)

The PLA assay detecting *in situ* protein interaction was performed using the kit (DUO92008, 92001, 92005, and 82049, Sigma-Aldrich). Purified ookinetes were fixed with 4% PFA for 30 min, permeabilized with 0.1% Triton X-100 for 10 min, and blocked with a blocking solution overnight at 4°C. The primary antibodies were diluted in the Duolink Antibody Diluent, added to the cells and then incubated in a humidity chamber overnight at 4°C. The primary antibodies were removed and the slides were washed with Wash Buffer A twice. The PLUS and MINUS PLA probe were diluted in Duolink Antibody Diluent, added to the cells and incubated in a pre-heated humidity chamber for 1 h at 37°C. Next, cells were washed with Wash Buffer A and incubated with the ligation solution for 30 min at 37°C. Then, cells were washed with Wash Buffer A twice and incubated with the amplification solution for 100 min at 37°C in the dark. Cells were washed with 1×Wash Buffer B twice and 0.01×Wash Buffer B once. Finally, cells were incubated with Hoechst 33342 and washed with PBS. Images were captured and processed using identical settings on a Zeiss LSM 880 confocal microscope.

### Protein immunoprecipitation

1.0×10^7^ ookinetes were lysed in 1 mL protein extraction buffer A plus (0.01% SDS, 1 mM DTT, 50 mM NaCl, 20 mM Tris-HCl, and pH8.0). After ultrasonication, the protein solution was incubated on ice for 15 min before centrifugation at 14,000g at 4 °C for 10 min. 1 μg of rabbit anti-HA antibody (#3724S, CST) and control IgG antibody (#2729, CST) were added to the supernatant respectively, and the solution was incubated on a vertical mixer at 4°C for 15 h. After incubation, 20 μl buffer A plus pre-balanced protein A/G beads (#20423, Pierce) was added and incubated for 5 h. The beads were washed three times with buffer A plus before elution with Laemmli buffer.

### Scanning electron microscopy analysis

The purified ookinetes were fixed in 2.5 % glutaraldehyde solution in 0.1 M phosphate buffer overnight, rinsed three times with PBS, and then post-fixed with 1% osmium tetroxide for 2 h. The fixed samples were dehydrated using a graded acetone series, CO2-dried in a critical-point drying device (K850, Emitech, USA) and gold-coated in a sputter coater (JFC-1600, JEOL, USA) as detailed previously [52]. The samples were imaged in a JSM-6390LV scanning electron microscope.

### Transmission electron microscopy analysis

Transmission electron microscope (TEM) experiments were performed using the protocol as described previously [53]. Purified ookinetes were pre-fixed with 4% glutaraldehyde in 0.1 M phosphate buffer at 4°C overnight, rinsed three times with PBS, post-fixed with 1% osmium acid for 2 h, and rinsed three times with PBS. The samples were dehydrated with concentration gradient acetone. After embedding and slicing, thin sections were stained with uranyl acetate and lead citrate prior to imaging. All samples were imaged under the HT-7800 electron microscope (USA).

### Peroxidase activity detection of mosquito midgut

Midgut staining with DAB (3,3-Diaminobenzidine) was performed as described [29]. Briefly, infected mosquito midguts were dissected 24 h post feeding. Midguts were fixed in 0.5% glutaraldehyde for 10 min at room temperature, washed in PBS, and developed for DAB activity. Samples were incubated at room temperature with 2.5 mM DAB (D8001, Sigma) and 1 mM H2O2 (18304, Sigma) in PBS (pH 6.5) and continuously observed under the microscope.

### NBD-Lipid Uptake

1-oleoyl-2-{6-[(7-nitro-2-1,3-benzoxadiazol-4-yl)amino]hexanoyl}-sn-glycero-3-phosphoserine (NBD-PS, #810194C), 1-oleoyl-2-{6-[(7-nitro-2-1,3-benzoxadiazol-4-yl)amino]hexanoyl}-sn-glycero-3-phosphoethanolamine (NBD-PE, #810155C), 1-oleoyl-2-{6-[(7-nitro-2-1,3-benzoxadiazol-4-yl)amino]hexanoyl}-sn-glycero-3-phosphocholin (NBD-PC, #810132C), and N-[6-[(7-nitro-2-1,3-benzoxadiazol-4-yl)amino]hexanoyl]-sphingosine-1-phosphocholine (NBD-SM, #810218C) were purchased from Avanti Polar Lipids (Birmingham, USA). NBD-phospholipid uptake experiments were performed according to the procedures [54]. 1.0×10^5^ ookinetes were collected from the *in vitro* culture, washed (2000 rpm for 5min at 22°C) and equilibrated at 22°C for 15 min in 500 μl of Hanks’ balanced salt solution (pH 7.4) containing 1g/ liter glucose (HBSS-glucose). A 500 μl of 3 mM NBD-phospholipid in HBSS-glucose was added to the ookinete suspension and incubated at 22°C for 1 h. After centrifugation (2000 rpm for 5min at 22°C) and discarding the supernatant, the ookinetes were mixed with 500 μl of ice-cold HBSS-glucose containing 5% fatty acid-free BSA (#9048-46-8, Sigma) to extract NBD-lipids incorporated into the exoplasmic leaflet of the ookinete plasma membrane, as well as unincorporated ones. The ookinete suspension was then placed in a petri dish for measuring the fluorescence signal of NBD-lipids inside the ookinetes. The fluorescence signals were captured using the Zeiss LSM 780 confocal microscope, and the signal intensity per cell was calculated using the ImageJ software. To test the effect of ATP depletion on the phospholipid uptake, ookinetes were pre-incubated in ATP depletion medium (HBSS without glucose but containing 20mM NaN3) for 1 h at 22°C before the NBD-lipid uptake assay [55].

### Phospholipid staining

For annexin V staining, the live ookinetes or dead ookinetes (after 60°C treatment for 5 min) were incubated in 1:100 Alexa488-Annexin V (#A13201, Invitrogen) and 1:100 propidium iodide (#P3566, Invitrogen) for 15 min, washed in PBS (2000 rpm at 22°C for 5min), and mounted on slides. For duramycin staining, the live or dead ookinetes were incubated in 0.25µg/mL biotinylated Duramycin [56] for 30 min, then stained with 1 µg/mL Alexa488-Streptavidin (#S32354, Invitrogen) and Hoechst 33342 for 30 min, washed in PBS (2000 rpm at 22°C for 5min), and mounted on slides. The ookinetes were analyzed using Zeiss LSM 780 confocal microscope.

### Auxin induced protein depletion

A stock solution of 250 mM Auxin/IAA (Indole 3-acetic acid, Sigma Aldrich, I2886) was prepared using the 100% EtOH. Mock treatment includes an equivalent volume of 100% EtOH. To determine the degradation efficiency of CDC50C-AID fusing protein, 1.0×10^5^ ookinetes were incubated with 1 mM IAA dissolved in ookinete media (RPMI 1640, 10% FCS, 100 μM XA, 25 mM HEPES, and pH 8.0) at 37°C for 1 h. After three times of wash in PBS (2000 rpm for 5min at room temperature), the cells were immediately fixed with 4% paraformaldehyde and transferred onto the slide for immunofluorescence assay. After imaging by Zeiss LSM 780 confocal microscope, the intensities of fluorescence signal of targeting proteins were quantitatively analyzed using the ImageJ software.

### Bioinformatics analysis and tools

The genomic sequences of *Plasmodium* genes were downloaded from the *Plasmodium* database of PlasmoDB (http://plasmodb.org). The genomic sequences of mosquito genes were downloaded from the *An. Stephensi* database (https://www.vectorbase.org/). The sgRNAs of target gene were designed using EuPaGDT (http://grna.ctegd.uga.edu/). The transmembrane domains of proteins were identified using the PROTTER Server (http://wlab.ethz.ch/protter/start/) [57]. Multiple sequence alignments were performed by ClustalW in MEGA7.0 [58].

### Quantification and statistical Analysis

Statistical analysis was performed using Graphpad Prism 8.0. Data collected as raw values are shown as mean±SEM or mean±SD. Two-tailed t-test or Mann-Whitney test was used to compare differences between treated groups and their paired controls. Details of statistical methods are reported in the figure legend. n represents the number of mosquitos or parasite cells tested in each group, or experimental replications.

---

**Figure S1.**
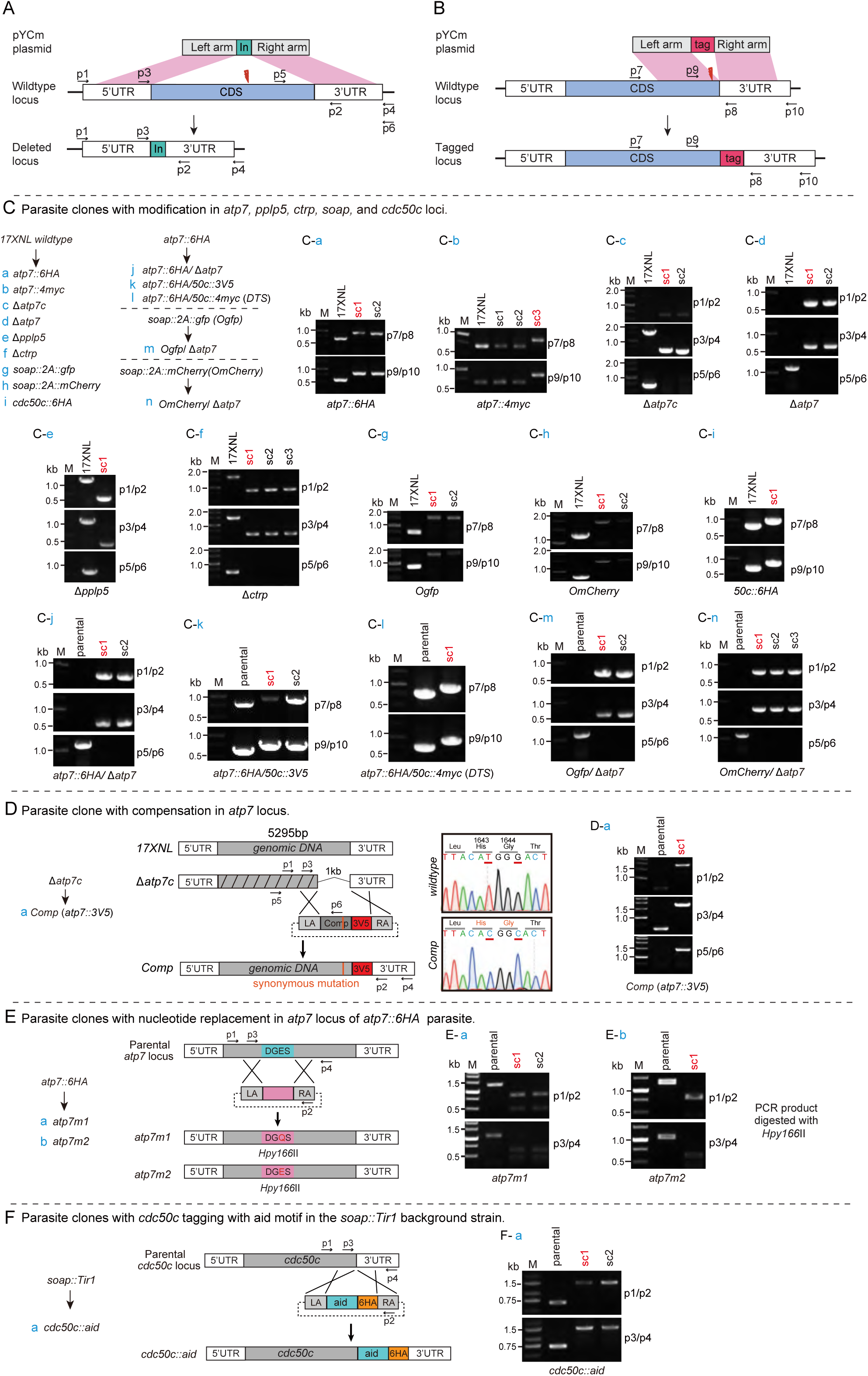
Genotyping of the modified parasites in this study. **A-B**. Schematic of CRISPR/Cas9-mediated gene editing via double cross homologous recombination, including gene deletion and gene tagging. **C.** For each modification, both 5’ and 3’ homologous recombination was detected using specific PCR pair (Supplementary Table 2) to confirm correct integration of the homologous template. Usually, one to three parasite clones (sc) for each modification were obtained after limiting dilution, and the clone indicated with red letter is used for phenotype and functional analysis. **D.** Diagram of ATP7 complementation in Δ*atp7c* mutant using CRISPR/Cas9 method. An epitope of triple V5 (3V5, red rectangle) was fused with the ATP7. Two synonymous mutations designed in the homologous template were confirmed via DNA sequencing. **E.** Diagram of residues substitutions in the endogenous ATP7 protein. From the parental *atp7::6HA* parasite, two mutant strains were generated: *atp7m1* with a missense substitution of E210Q, and *atp7m2* with a synonymous substitution of E210E. **F.** Diagram of tagging endogenous *cdc50c* with an *aid::6HA* motif for auxin-induced protein degradation. In the parental parasite *soap::Tir1*, endogenous *cdc50c* was tagged with *aid::6HA* tag using the CRISPR/Cas9 method.

**Figure S2.**
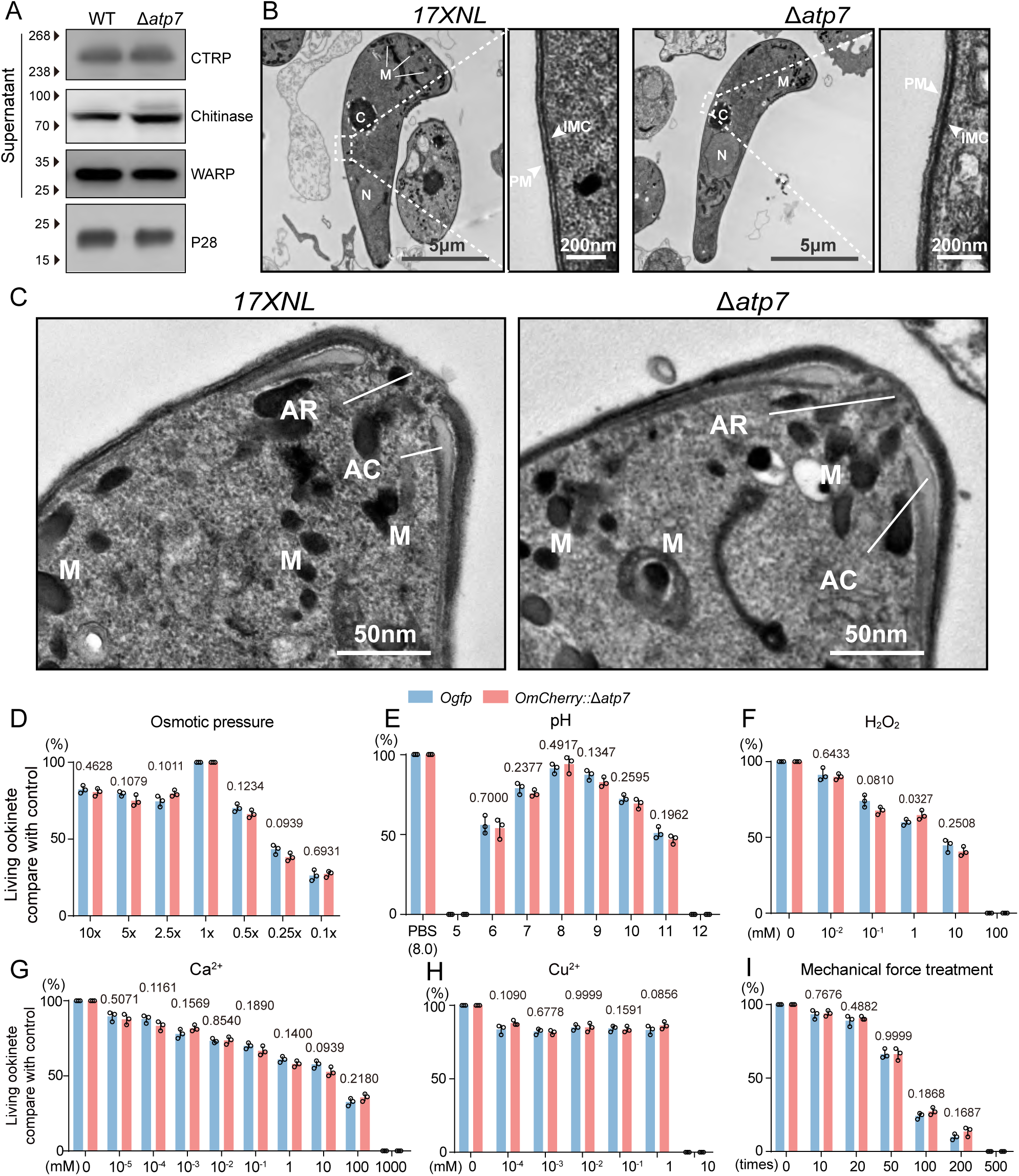
Wildtype and ATP7-depleted ookinete microneme secretion, apical structure, and parasite survival to stress. **A.** Immunoblot of microneme-secreted proteins (CRTP, Chitinase, and WARP) by WT and Δ*atp7* ookinete cultures. Plasma membrane protein P28 was used as a loading control. **B.** Transmission electron microscope (TEM) analysis of WT and *Δatp7* ookinetes. M: microneme, N: nuclear, IMC: inner membrane complex, and C: crystalloid body. **C.** Enlarged TEM images showing ookinete apical structures. AR: apical ring, AC: apical collar, M: microneme. **D-I**. Morphology and survival analysis of ATP7-deficient ookinetes under stress conditions *in vitro*. Two reporter strains *Ogfp* and *OmCherry*/Δ*atp7* were generated and tested in this experiment (Fig. S3). Equal number of *Ogfp* (GFP) and *OmCherry*/Δ*atp7* (mCherry) mature ookinetes were mixed and incubated in PBS with different artificial stress factors, including osmotic pressure (**D**), pH (**E**), peroxide (**F**), Ca^2+^ (**G**), Cu^2+^ (**H**) and mechanical force (**I**). Ookinete morphology and survival was analyzed after 5 h incubation. Different PBS dilutions were used in **D**, different pH in **E**, and different molecule or ions concentrations in **F**, **G** and **H**. In **I**, the ookinetes were extruded from a sterile syringe (26G) at different rates. The percentage of morphologically normal ookinetes was normalized to that of the control without stress. Data are shown as mean ± SEM and analyzed by Two-tailed unpaired Student’s t test. All experiments were repeated three times.

**Figure S3.**
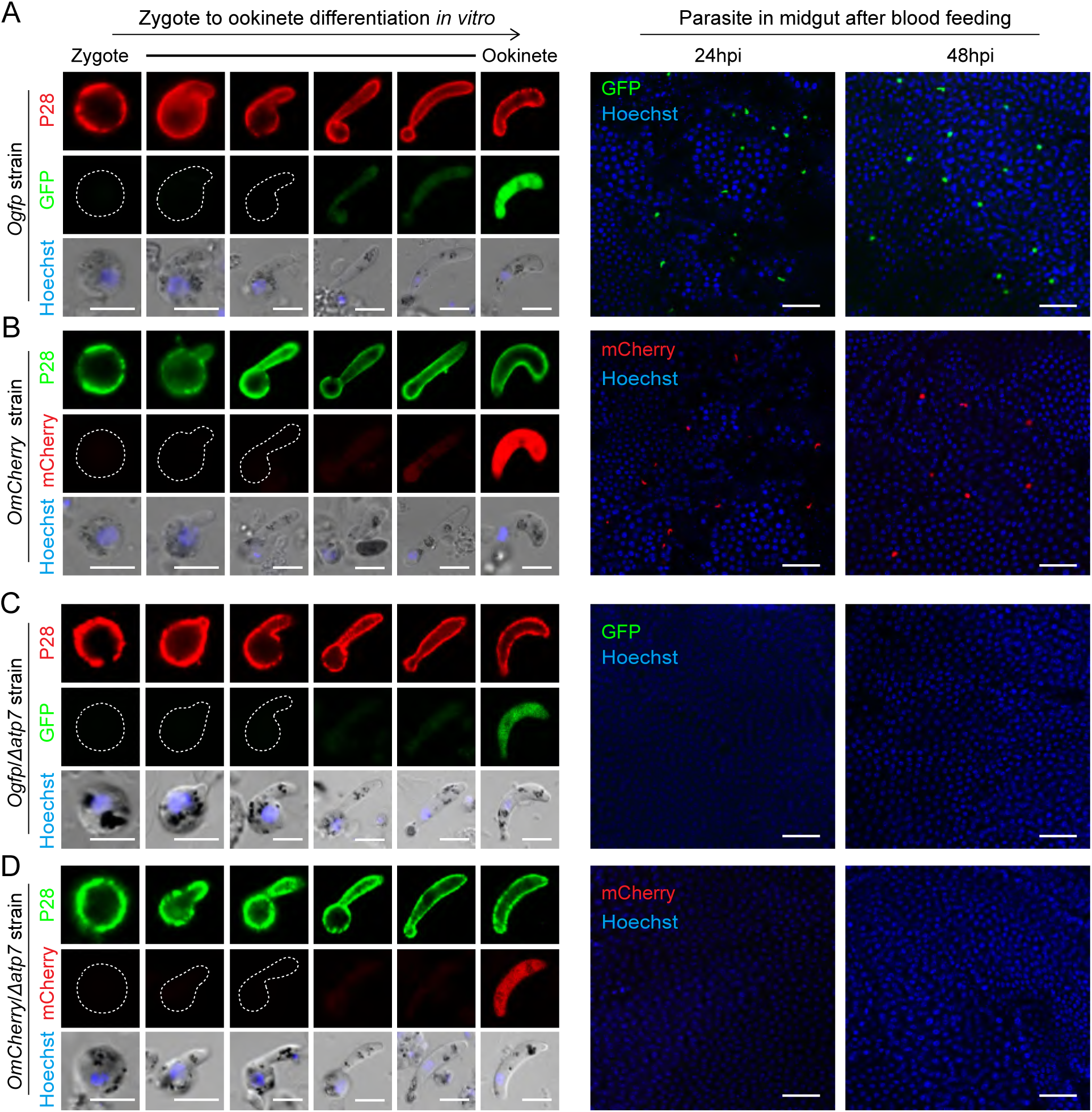
Characterization of four transgenic strains expressing GFP or mCherry in ookinetes. Fluorescent protein sequences (*gfp* and *mCherry*) were fused to the endogenous gene *soap*, an abundant ookinete and early oocyst protein, generating the *Ogfp* strain and *OmCherry* strain (see Fig S1C). GFP expression in the *Ogfp* strain (**A**) and mCherry expression in the *OmCherry* strain (**B**) from zygote to ookinete differentiation (left panel) and in the midgut of infected mosquitoes at 24 and 48 h pi (right panel). The *atp7* gene was disrupted in the *Ogfp* and *OmCherry* strains, generating the *Ogfp*/Δ*atp7* (**C**) and *OmCherry*/Δ*atp7* (**D**) strains, respectively. P28 is an ookinete plasma membrane protein. Nuclei are labeled with Hoechst 33342. Scale bar is 5 μm in left panels and 50 μm in right panels.

**Figure S4.**
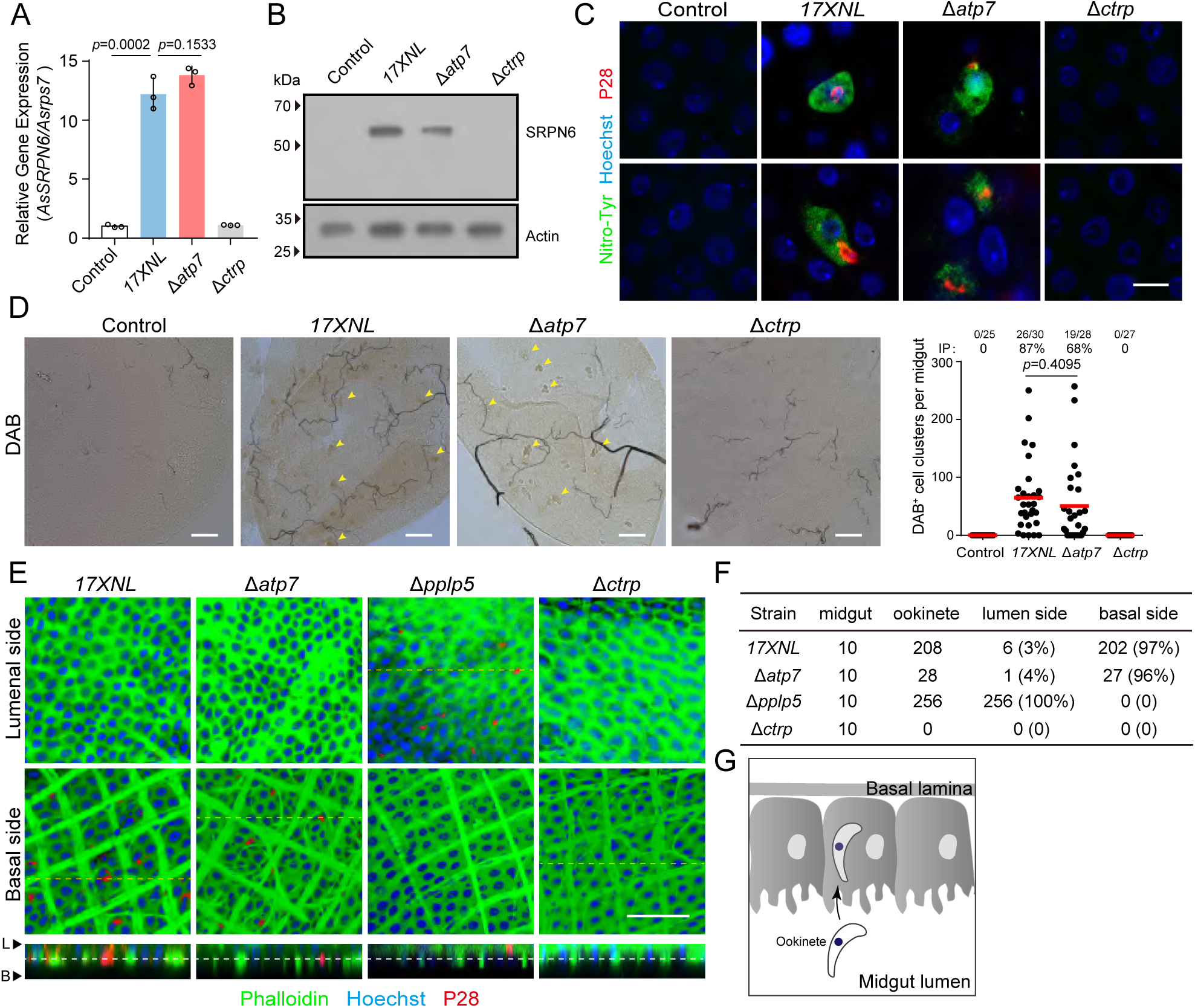
ATP7-depleted ookinetes are able to invade the midgut epithelium. **A.** qRT-PCR of the *Anopheles stephensi* midgut *SRPN6* mRNA relative to the housekeeping gene *Asrps7* (24 h pi) from the naïve blood (control)-fed mosquitoes or parasite (17XNL, △*atp7*, and △*ctrp*)-infected mosquitoes. **B.** Immunoblot of the SRPN6 protein in midguts at 24 h pi. Actin was used as a loading control. **C.** Staining of infected midguts for nitrotyrosine and P28 at 24 h pi. P28 is an ookinete plasma membrane protein. Epithelia cells invaded by ookinetes are positive for nitrotyrosine. **D.** Peroxidase activity of infected midguts via DAB assay on 24 h pi. Yellow arrows point to DAB^+^ signals. The right panel shows quantification of the results. **E.** 3D-scan positional analysis of ookinetes in mosquito midguts using confocal microscopy. Parasite-infected midguts were dissected at 18 h pi and visualized after staining with parasite P28 antibody, phalloidin (for mosquito actin), and Hoechst 33342. Thick muscle actin filaments of mosquito are observed in the midgut basal sides. **F.** Quantification of results in **E**. Ten midguts were measured for each group. **G.** Diagram showing successful epithelium invasion of the ATP7-depleted ookinetes. Scale bar is 10 μm in **C**, and 50 μm in **D** and **E**. Data are shown as mean ± SD in **A**. In **D**, the numbers on the top are the number of DAB^+^ midguts / the number of midgut measured; IP: infection prevalence. Red horizontal lines show mean values. Data were analyzed by two-tailed unpaired Student’s t test in **A** and Mann–Whitney test in **D**. All experiments in this figure were independently repeated three times.

**Figure S5.**
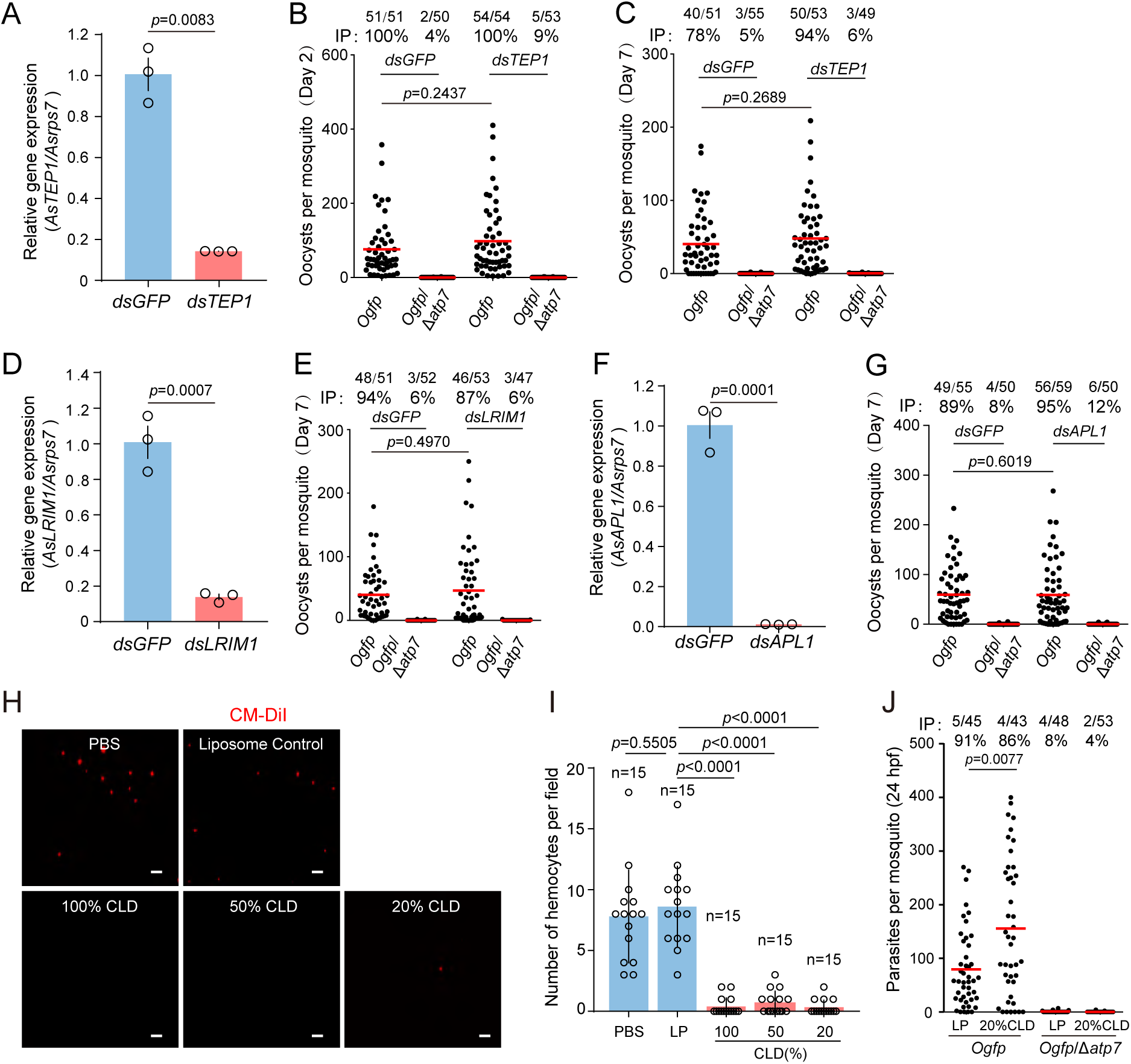
Elimination of Δ*atp7* ookinetes is independent from the mosquito complement immunity. **A.** TEP1 mRNA abundance in mosquitoes 24 h after injection with TEP1 dsRNA (*dsTEP1*) compared with controls injected with GFP dsRNA (*dsGFP*). Expression level of *TEP1* was normalized to the housekeeping gene *Asrps7*. **B-C**. Midgut oocyst development of *Ogfp* and *Ogfp*/Δ*atp7* in mosquitoes treated with *dsGFP* or *dsTEP1*. Mosquitoes were dissected at 2 (**B**) and 7 days (**C**) post infection. **D.** LRIM1 mRNA abundance in mosquitoes 24 h after injection with LRIM1 dsRNA. **E.** Midgut oocyst development of *Ogfp* and *Ogfp*/Δ*atp7* in dsRNA injected mosquitoes. **F.** APL1 mRNA abundance in mosquitoes 24 h after injection with APL1 dsRNA. **G.** Midgut oocyst development of *Ogfp* and *Ogfp*/Δ*atp7* in dsRNA injected mosquitoes. **H.** Detection and depletion of mosquito hemocytes. Hemocytes were stained with CM-DiI, a lipophilic dye specifically labeling mosquito hemocytes. Different dilutions of clodronate liposome (CLD) were injected into mosquito hemocoel for hemocyte depletion. Liposome was used as a control. Scale bar: 50 μm. **I.** Quantification of mosquito hemocyte depletion in **H**. **J.** Midgut oocyst development of *Ogfp* and *Ogfp*/Δ*atp7* in mosquitoes with hemocytes depleted by 20% CLD. Mosquitoes were dissected at 24 h post infection. Data are shown as mean ± SD in **A**, **D**, **F**, and **I**. In **B**, **C**, **E**, **G**, and **J**, the numbers on the top are the number of midguts containing parasites / the number of midguts measured; IP: infection prevalence. Red horizontal lines show the mean value of cell numbers. Data were analyzed by two-tailed unpaired Student’s t test in **A**, **D**, **F**, and **I**, and **J** and Mann–Whitney test in **B**, **C**, **E**, **G**, and **J**. All experiments in this figure were independently repeated three times.

**Figure S6.**
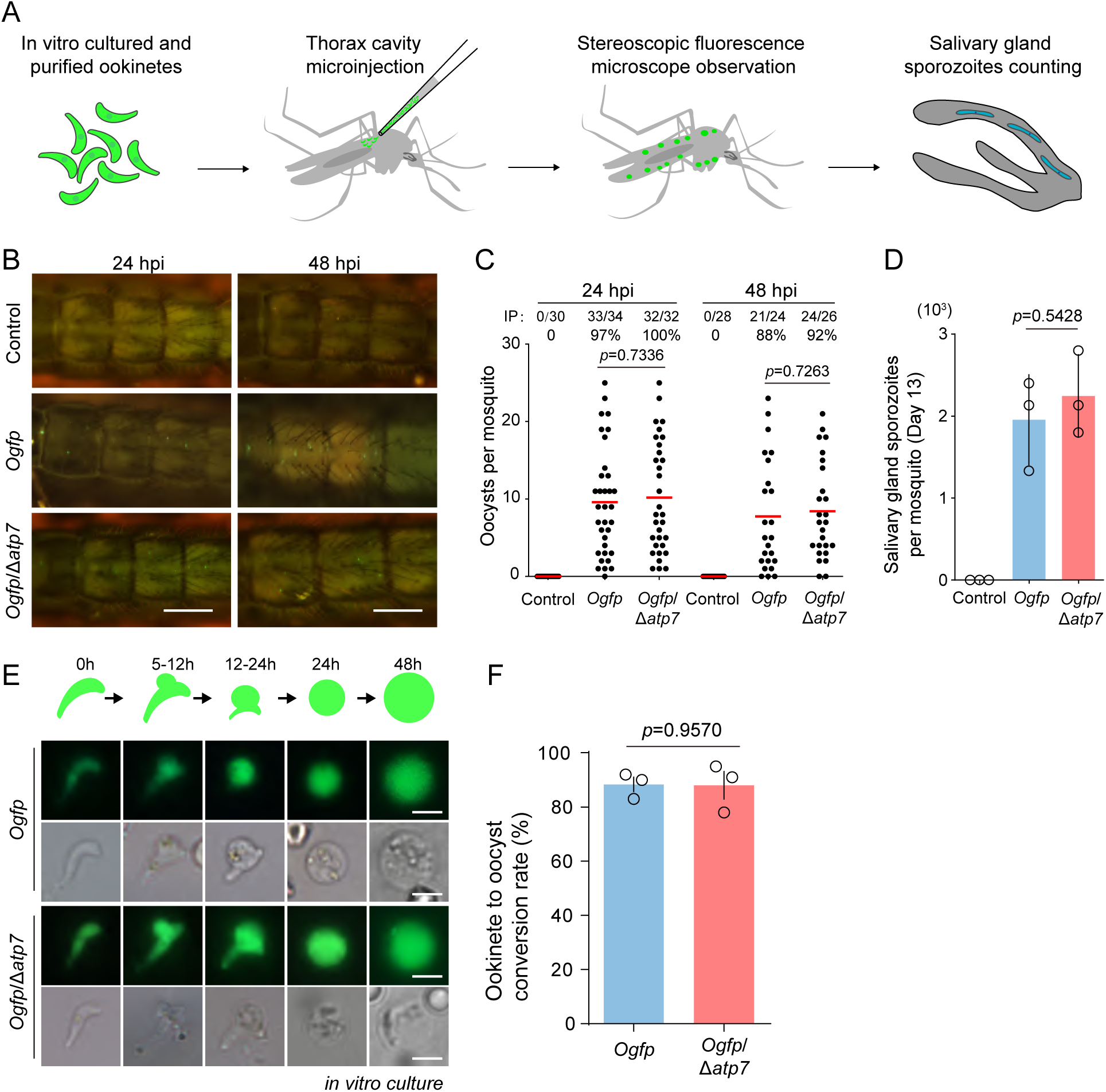
Ookinete microinjection into the mosquito hemocoel restores the Δ*atp7* defects. **A.** Schematic of mosquito injection of *in vitro* cultured ookinetes into the hemocoel. Approximately 690 *in vitro* cultured *Ogfp* and *Ogfp*/Δ*atp7* ookinetes were injected per mosquito through thorax cavity. Hemocoel oocyst and salivary gland sporozoite number were determined. **B.** Images of GFP^+^ oocysts in mosquito hemocoel at 24 and 48 h post ookinete injection. PBS was injected as a control. Scale bar=500 μm. **C.** Quantification of GFP^+^ oocyst numbers in **B**. **D.** Salivary gland sporozoite numbers at day 13 post ookinete injection. **E.** Analysis of *in vitro* ookinete to early oocyst transformation of *Ogfp* and *Ogfp*/*Δatp7* parasites. Scale bar=5 μm. Upper diagrams indicate the morphological changes from ookinete to early oocyst *in vitro*. **F.** Quantification of the *in vitro* ookinete to early oocyst transformation in **E**. Data are shown as mean ± SEM in **D** and **F**. In **C**, the numbers on the top are the number of mosquitoes carrying oocysts / the number of mosquitoes observed; IP: infection prevalence; red horizontal lines show the mean value of oocyst numbers. Two-tailed unpaired Student’s t test in **D** and **F**, and Mann–Whitney test in **C**. All experiments in this figure were independently repeated three times.

**Figure S7.**
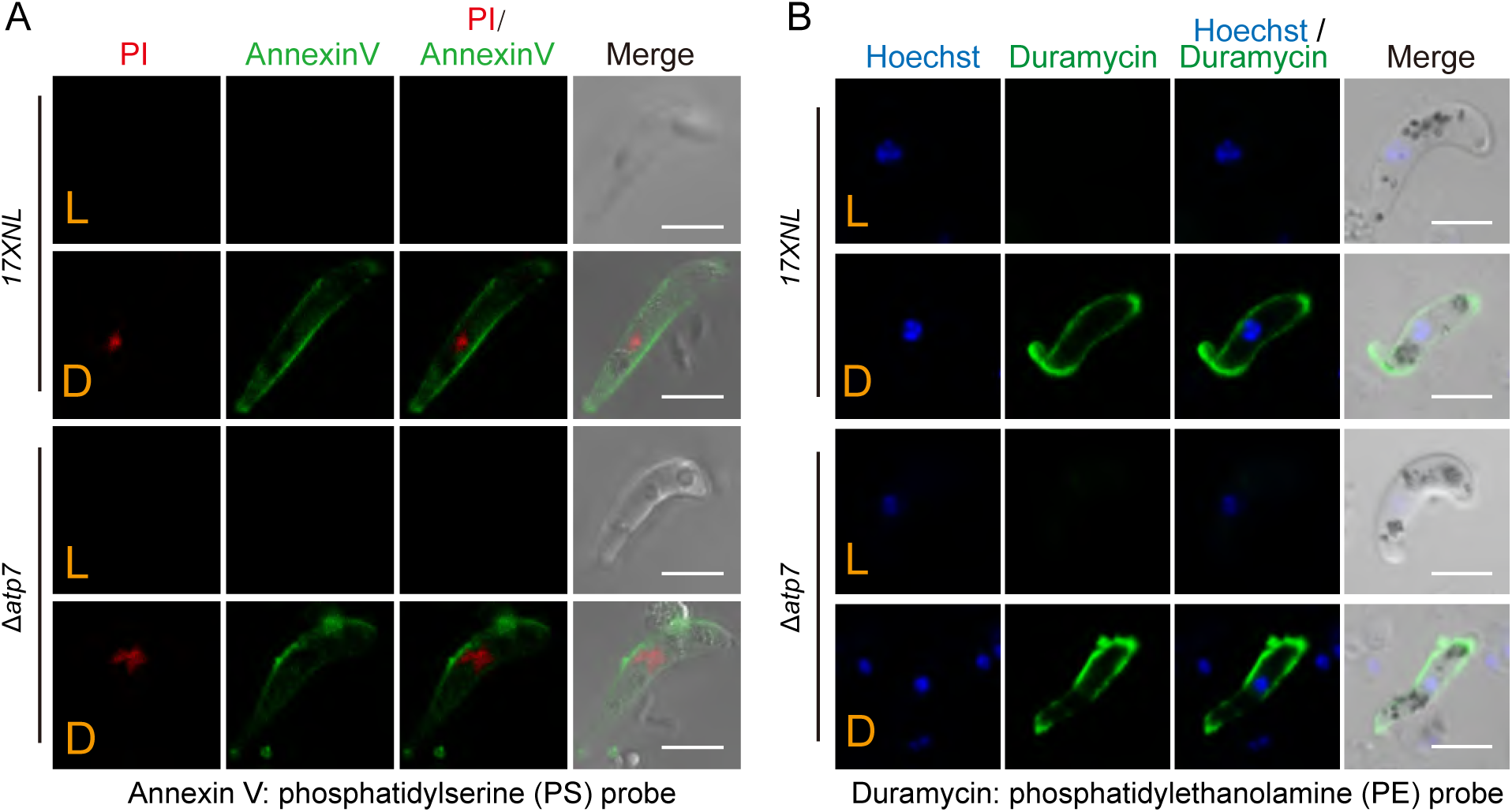
Detection of cell-surface phospholipids (PS and PE) in WT and Δ*atp7* ookinetes. **A.** Double staining of 17XNL and Δ*atp7* ookinetes with Annexin V (PS probe) and PI (propidium iodide, cell death marker). Scale bar is 5 μm. **B.** Double staining of the 17XNL and Δ*atp7* ookinetes with Duramycin (PE probe) and Hoechst 33342 (nuclear stain). Scale bar is 5 μm. L, live ookinetes; D, dead ookinetes killed by treatment at 60°C for 5 min.

**Figure S8.**
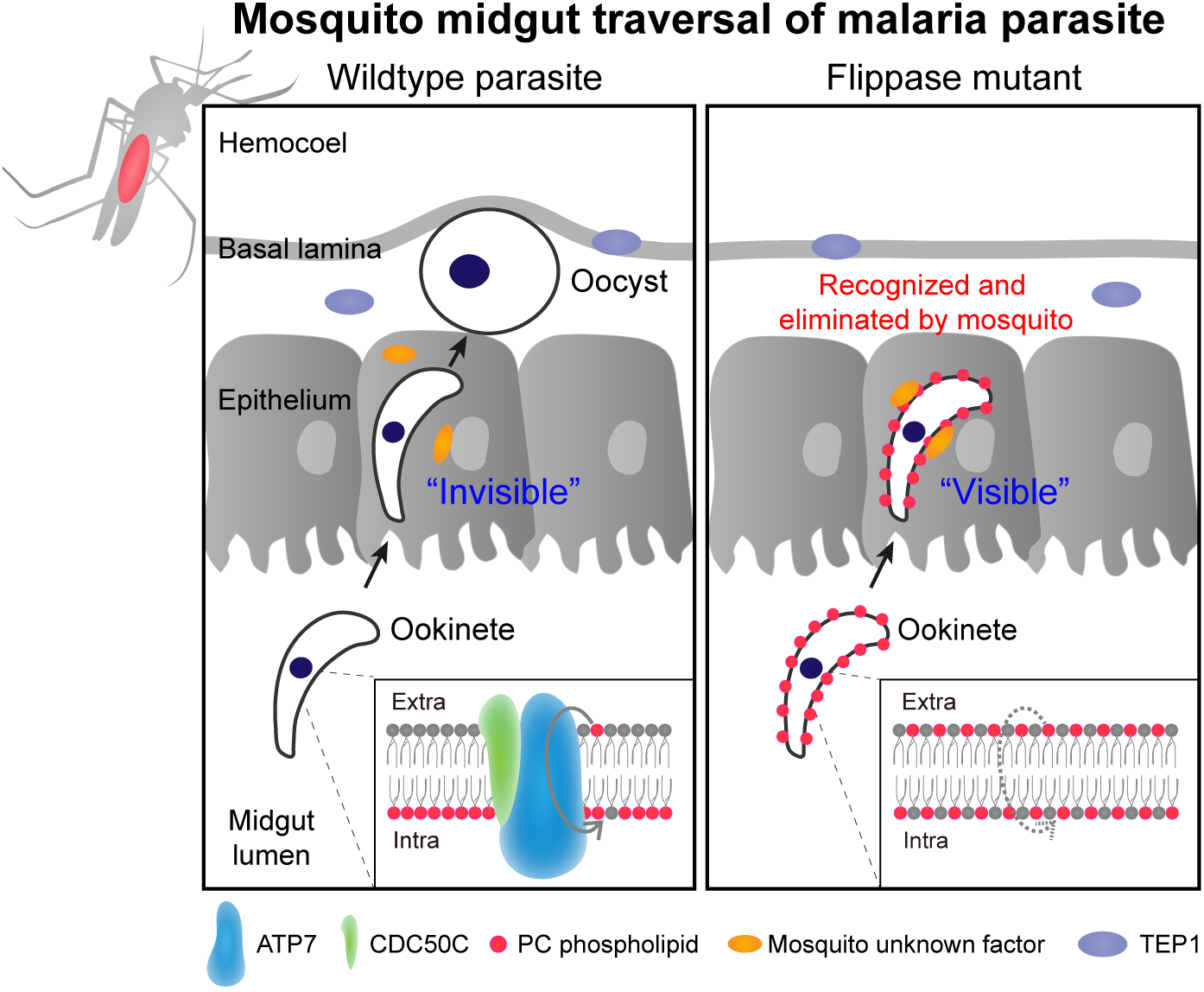
Proposed model of ATP7/CDC50C transporting phospholipid PC from theouter leaflet to the inner leaflet of ookinete plasma membrane, which likely enable ookinete “invisible” and evade mosquito immune recognizing and attack inside the midgut epithelium.

